# Spatial control of cell division by GA-OsGRF7/8 module in a leaf explains the leaf length variation between cultivated and wild rice

**DOI:** 10.1101/2021.05.06.443003

**Authors:** Vikram Jathar, Kumud Saini, Ashish Chauhan, Ruchi Rani, Yasunori Ichihashi, Aashish Ranjan

## Abstract

- Cellular and genetic understanding of rice leaf size regulation is limited, despite rice being the staple food of more than half of the global population. We investigated the mechanism controlling the rice leaf length using cultivated and wild rice accessions that remarkably differed for leaf size.
- Comparative transcriptomics, Gibberellic Acid (GA) quantification, and leaf kinematics of the contrasting accessions suggested the involvement of GA, cell cycle, and Growth-Regulating Factors (GRFs) in the rice leaf size regulation. Zone-specific expression analysis and VIGS established the functions of specific GRFs in the process.
- The leaf length of the selected accessions was strongly correlated with GA levels. Higher GA content in wild rice accessions with longer leaves and GA-induced increase in the leaf length via an increase in cell division confirmed a GA-mediated regulation of division zone in rice. Downstream to GA, *OsGRF7* and *OsGRF8* function for controlling cell division to determine the rice leaf length.
- Spatial control of cell division to determine the division zone size mediated by GA and downstream OsGRF7 and OsGRF8 explains the leaf length differences between the cultivated and wild rice. This mechanism to control rice leaf length might have contributed to optimizing leaf size during domestication.

## Introduction

The basic leaf development program includes leaf primordia initiation on the flanks of shoot apical meristem that follows growth and patterning by multiple transcription factor families (Townsley & Sinha, 2012; Bar & Ori, 2014; Du *et al*., 2018). While leaf initiation and patterning are regulated by a suite of transcription factors and complex hormonal interactions, the final leaf size is determined by the coordination of cell division and expansion (Li *et al*., 2007; Gonzalez *et al*., 2012; Chitwood & Sinha 2016; Sarvepalli *et al*., 2019). Considering the leaf as the major plant organ for light-harvesting and perception of environmental signals, understanding the genetic basis of leaf size regulation is imperative for crop improvement.

The regulation of leaf size is extensively studied for simple leaves of Arabidopsis and compound leaves of tomato, however detailed mechanistic studies are relatively scarce for monocot cereals except for maize (Efroni *et al*., 2010; Rodriguez *et al*., 2013; Vercruysse *et al*., 2020a). The spatial and temporal regulation of cell proliferation and cell elongation to determine the final leaf size varies between monocots and dicots. Cell proliferation and expansion are temporally separated in dicots as cell proliferation predominates in young developing leaf, thereafter cells expand before reaching maturity (Beemster *et al*., 2005; Piazza *et al*., 2005; Andriankaja *et al*., 2012). In contrast, cell proliferation and expansion are spatially separated in monocots, such as maize, where division, elongation, and maturation zones are present together at steady-state leaf growth (Tardieu & Garnier, 2000; Granier & Tardieu, 2009; Fina *et al*., 2017). Monocot leaves with such spatial resolution of the growth zones allow the systematic study of the zones and their contribution in the leaf size control.

Changes in the size of the division and/or elongation zone alter the final leaf size in Arabidopsis and maize (Tardieu & Granier, 2000; Tardieu *et al*., 2000; Gonzalez *et al*., 2010; Kalve *et al*., 2014; Gazquez & Beemster, 2017; Nelissen *et al*., 2018). Several phytohormones play important roles in the regulation of cell division, elongation, and transition between division and elongation (Inze & De Veylder, 2006; Su *et al.*. 2011; Avramova *et al*., 2015; Takatsuka & Umeda, 2014). GA is classically known for mediating cell-elongation response, however recent studies in Arabidopsis and maize have shown its key roles in determining cell division (Achard *et al*., 2009; Nelissen *et al*., 2012). GA, after being perceived by a receptor protein GIBBERELLIN INSENSITIVE DWARF1 (GID1), promotes degradation of DELLA repressor proteins as well as suppresses expression of DELLA-encoding *SLR1* in feedback regulation (Willige *et al*., 2007; Davière & Achard, 2013; Shan *et al*., 2014). GA-mediated suppression of DELLA proteins results in activation of GA-responsive transcription factors, such as GROWTH-REGULATING FACTORs (GRFs) (Omidbakhshfard *et al*., 2015). GRFs are reported to directly regulate cyclins to control cell division in different plant species (van der Knaap *et al*., 2000; Choi *et al*., 2004; Nelissen *et al*., 2015; Fina *et al*., 2017). Arabidopsis plants overexpressing *AtGRF1, AtGRF2*, and *AtGRF3* develop longer leaves and larger organ sizes, whereas *grf* mutants produce smaller and narrower leaves (Kim *et al*., 2003; Beltramino *et al*., 2018). Moreover, *GRF* expression modulation in maize alters leaf length with associated changes in division zone size (Wu *et al*., 2014; Nelissen *et al*., 2015)

A detailed cellular basis and underlying genetic and hormonal regulation of rice leaf length control are not systematically investigated. Rice domestication has significantly reduced the genetic variability in the cultivated varieties (Huang *et al*., 2012; Chen *et al*., 2021). Therefore, a comparative study of leaf size and associated cellular differences across cultivated and wild rice would provide a comprehensive genetic understanding of the rice leaf size regulation. Several wild relatives of rice grow taller and accumulate higher biomass associated with larger organ size including leaf size (Sanchez *et al*., 2013; Mathan *et al*., 2021a). GA is known to regulate stem elongation in rice, and manipulation of GA-biosynthesis and signaling genes have optimized plant height during the green revolution (van der Knaap *et al*., 2000). Several GRFs downstream to GA are known to regulate various aspects of rice development and architecture (Luo *et al*., 2005; Kuijt *et al*., 2014; Gao *et al*., 2015; Tang *et al*., 2018; Chen *et al*., 2020). OsGRF1, the founding member of the rice GRF transcription factor family, is expressed in intercalary meristems of internodes and influences rice leaf growth (van der Knaap *et al*., 2000; Lu *et al*., 2020). However, the contribution of cell division and its regulation by GA and specific downstream GRFs for rice leaf length regulation is not systematically elucidated. Further, the signaling module that explains the leaf length differences between the contrasting cultivated and wild rice is also not investigated. Our study has systematically integrated GA with the rice leaf length control via cell cycle regulation and identified specific GRFs, OsGRF7 and OsGRF8, downstream to GA that not only explains the leaf length differences between the selected cultivated and wild rice but also would provide a way to optimize rice leaf length for improved physiological performance.

## Materials and Methods

### Plant materials and growth conditions

The genotypes used for this study are summarized in Table S1. Rice seeds were first germinated on Petri plates containing wet germination paper, followed by transfer of seedlings to a hydroponic solution containing one-fourth MS Media (Himedia) in standard concentrations. The nutrient solution was changed every third day during the seedling growth. Seedlings were grown at 28 °C day and 23 °C night temperature with 14:10 hours of light and dark cycle. For exogenous treatments, 10μM of gibberellic acid (GA_3_), and 1μM of gibberellic acid biosynthesis inhibitor Paclobutrazol (PAC) were added in hydroponic solution.

### Phenotyping of leaf morphological traits and leaf growth analysis

The quantification of leaf length and width was performed on each fully-grown leaf from the second to eight leaves of all the selected rice accessions. Leaf length and width were quantified manually using a ruler. Leaf-sheath length was measured as a distance between the seedling base to ligule. The leaf-blade length was quantified as the distance between the ligule to the leaf tip. Leaf-blade width was measured in the central widest part of the fully expanded leaf blade. Total mature leaf length was calculated by adding leaf sheath and leaf blade length. The leaf growth analysis was done on the fourth leaf, when the leaf emerged from the sheath of the third leaf, as explained by Sprangers *et al*. (2016). Leaf length was measured daily from the day of emergence up to the seventh day by considering the seedling base as the starting point of the leaf. Leaf elongation rate (LER) was calculated as the leaf growth in one day divided by the time interval.

### RNA-seq library preparation and sequencing

The fourth leaves of the selected accessions were collected on the second day after leaf emergence for RNA-seq library preparation as described by Sprangers *et al*., 2016. Leaves were in the primordia 5 (P5) stage and were growing at a steady growth rate at the time of collection. Libraries were prepared in three biological replicates using YourSeq Full Transcript RNAseq Library Kit (AMARYLLIS NUCLEICS) as per the manufacturer’s protocol. These RNA-seq libraries were sequenced on the Illumina HiSeq platform, and reads were generated in 150bp paired-end format.

### Quality filtering, mapping of reads, and differential gene expression analysis

The reads were quality filtered using the NGSQC toolkit that involved the removal of adapter sequences and low-quality reads (Phred score <30) (Patel *et al*., 2012). The quality-filtered reads were mapped to reference cDNA sequences of rice from the Rice Genome Annotation Project database (RGAP 7) using default parameters of bowtie2 (Langmead & Salzberg, 2012). SAMtools were used to generate bam alignment files that were used to tabulate raw counts for each gene (Li *et al*., 2009). The raw counts were normalized to library size and corrected for library composition bias using the Trimmed Mean of M values normalization approach (Robinson and Oshlack, 2010). edgeR was used for differential gene expression analysis (Robinson *et al*., 2010). glm approach with quasi-likelihood F-test was used for differential gene expression analysis. Significant differentially expressed genes for each pair-wise comparison were determined based on FDR cut-off < 0.05 and logFC value >1 and <-1.

### Principle Component Analysis with Self Organizing Map (PCA-SOM) clustering

Normalized read counts (Dataset S7) were used for a gene expression clustering method (Chitwood *et al*., 2013). For clustering, the genes in the upper 50% of the coefficient of variation for expression across different accessions were used. The scaled expression values were used to generate accession-specific gene clusters (Wehrens & Buydens, 2007). Three by three hexagonal SOM clusters and 100 training interactions were used during clustering. The SOM clusters were visualized in PCA space, where PC values were calculated based on gene expression across the accessions with the help of the prcomp R package. The heatmaps were generated for each cluster using superheat R package.

### GO Enrichment Analysis

GO enrichment analysis was performed using the Plant Transcriptional Regulatory Map GO enrichment tool of the Plant Transcription Factor Database (Tian *et al*., 2019).

### Gene regulatory network analysis

An unsigned correlation network was constructed using the weighted gene correlation network (WGCNA) package version 1.68 (Langfelder *et al*., 2008). Data were z-score normalized (R scale function). The soft-thresholding power was chosen based on a scale-free topology with a fit index of 0.8. An adjacency matrix with the selected soft-thresholding power was calculated and transformed into a topological overlap matrix (TOM). Using the TOM, network properties such as strength were calculated and the network was visualized using the igraph package version 1.2.5 (Csardi *et al*., 2006) and visNetwork 2.0.9 (https://datastorm-open.github.io/visNetwork/). A Fast-Greedy modularity optimization algorithm was selected to define modules in the integrated network.

### Leaf kinematics analysis

To estimate the division zone size, the basal 4 cm segment of the fourth leaf was collected at different days of emergence up to day five as described by Sprangers *et al*. (2016). Samples were placed in 3:1 (v/v) absolute ethanol: acetic acid, rinsed for 30 min in a buffer containing 50 mM NaCl, 5 mM EDTA, and 10 mM Tris-HCl, pH 7, and nuclei were stained by incubating the samples in the dark for 2 min in the same buffer solution containing 1 mg/ml 49,6-diamino-phenylindole (DAPI). Fluorescent nuclei were observed with a microscope with an epi-fluorescent condenser (Nikon 80i-epifluorescence Microscope), and the images obtained were used to measure the division zone size as the distance between the base of the leaf and the most distal mitotic cell. To determine cell length profiles, fourth leaves at the P5 developmental stage were collected on the second day after leaf emergence at the steady-state growth. The leaves were kept in absolute ethanol for 48 h (with one change at 24 h), then transferred to lactic acid, and stored at 4°C. Samples were mounted on microscope slides using glycerol. The epidermal cells at each millimeter length were visualized using a phase-contrast microscope (LMI, UK) at 40X magnification. The various growth parameters from the leaf kinematics experiments were quantified using formulae explained in Sprangers *et al*. (2016).

### GA quantification

The fourth leaves at the P5 developmental stage were collected on the second day after leaf emergence at the steady-state growth, similar to leaf kinematics and transcriptomics experiments, for GA quantification in full leaves. For zone-specific GA quantification, the samples were collected from two different zones of fourth leaves. Zone I was up to 0.8 cm from the leaf base and zone II was from 0.8 cm to 1.6 cm. The samples were lyophilized for 72 hr and finely ground in liquid nitrogen. 25 mg of each sample was extracted in ice-cold buffer (MeOH: H2O: HCOOH, 15:4:0.1) on a shaker for overnight at 4°C. The internal standard was added to the extract and vortexed for 10 min. The extract was centrifuged for 15 min at 16,000 rcf at 4°C, the supernatant was passed onto a pre-conditioned (MeOH and 0.1% of HCOOH) C18 RP SPE column, and the column was washed with 0.1% HCOOH and 5% MeOH. Finally, elution was done with ice-cold 0.1% HCOOH in Acetonitrile. After evaporation of the solvent, the pellet was re-suspended in 5% MeOH. GA content was then analyzed using LC-MS/MS.

### RNA isolation, cDNA synthesis, and qRT-PCR analysis

For zone-specific expression profiling, samples from zone I and II of the fourth leaves were collected as explained above for zone-specific GA quantification. Total RNA was isolated using Trizol reagent (Invitrogen) and treated with RNase-free DNase I (Thermo Fisher Scientific). 1.5 μg of total RNA was converted to first-strand cDNA using RevertAid First-Strand cDNA Synthesis Kit (Thermo Fisher Scientific). The quantitative real-time-PCR analysis was performed on the CFX Connect Real-Time System (Bio-rad) using PowerUp^™^ SYBR^®^ Green fluorescence dye (Thermo Fisher Scientific). The primer pairs for qRT-PCR were designed based on gene sequences obtained from RGAP 7 (Table S6). The primer efficiency was analyzed using a standard curve method. From each species five different cDNA concentration (25,50,100,200 and 400ng) used to generate standard curve and primer efficiency was calculated using the formula E = (10[−1/slope] 257 −1) *100. The primer specificity of amplified gene products across the selected rice genotypes was viewed through dissociation curve analysis. Relative expressions of the target genes were analyzed using actin (LOC_Os03g50885.1) as the internal control applying the 2^-ΔΔCt^ method for fold change and 2^-ʔCt^ method for relative transcript levels quantification (Livak & Schmittgen, 2001). The expression pattern of each gene was confirmed by three independent biological experiments.

### Transient gene silencing line generation

Virus-induced gene silencing (VIGS) mediated transient silencing lines for *OsGA20OX2*, *OsSLR1*, *OsGRF7*, and *OsGRF8* genes were generated using the mVIGS vector as previously described by Kant et al. (2017). Recombinant vector constructs were transformed into Agrobacterium strain EHA105, and a single Agrobacterium colony was used for primary culture. After overnight incubation at 28 °C, a secondary culture was grown to an OD600 of 0.6 to 0.8 with 200 μM acetosyringone. Cells were harvested after centrifugation and resuspended in 10 mM MES, 10 mM MgCl2, and 200 μM acetosyringone. The meristematic region located at the basal region (junction of shoot and root) of 6-day old rice seedlings were injected with 50 μL of bacterial suspension using a vertically positioned syringe as described by Kant et al., 2015. After inoculation seedlings were transferred onto sterile Whatman No. 1 filter paper immersed in MS medium placed on a solid support with its ends dipped into a reservoir containing the nutrient medium. To avoid drying, plant roots were covered with moist tissue paper and transferred to tubes containing MS medium 24 h post-inoculation and were maintained at 28°C under conditions described above. As positive controls, we generated transient silencing lines for chlorophyll biosynthesis gene (*ChlH*) and confirmed positive transgenic plants by observing leaves’ chlorosis phenotype and gene expression analysis (Fig. S9a-d).

## Results

### Leaf size variation across the selected cultivated and wild rice accessions

Quantification of the leaf length and the maximum leaf blade width of the fully-grown second to eighth leaves of the twelve rice accessions showed a strong variation in leaf length and width (Tables 1, S2). The length of the fully-grown eighth leaf varied from 44.8 cm in a cultivated rice *Oryza sativa ssp. indica* cv. IR64 to 87.8 cm in a wild rice *Oryza australiensis* (Table 1). The length of the fully-grown eighth leaf was significantly higher in the five wild rice accessions *O. australiensis, O. rufipogon, O. latifolia, O. glumaepatula*, and *O. punctata* than the Asian cultivated rice varieties and the African cultivated rice *O. glaberrima*. Significant variation in leaf width was also observed across the cultivated and wild rice accessions (Table S2). We then selected four genetically diverse accessions, which captured the leaf length variations across the investigated accessions, for detailed phenotypic and genetic characterization. The four selected accessions were: IR64 (an *indica* variety of the cultivated species *O. sativa)* and Nipponbare (a *japonica* variety of the cultivated species *O. sativa*) with smaller leaves, the wild rice *O. australiensis* with the longest leaves, and *O. glaberrima* (the African cultivated rice) with intermediate leaf length (Table 1; Fig. 1a). The leaf phenotype of second to tenth leaves of the four accessions further confirmed the variation in leaf length pattern (Fig. 1b, S1).

**Table 1.** Quantification of leaf length (cm) of fully-developed second leaf to the eighth leaf of the seven cultivated and five wild rice accessions. Each value represents mean ± SD, where n = 15 data points from different plants. Different letters indicate statistically significant differences according to one-way ANOVA followed by post-hoc Tukey HSD calculation at P < 0.05.

| Oryza species | Cultivar/<br>accession<br>number | Leaf number |  |  |  |  |  |  |
| --- | --- | --- | --- | --- | --- | --- | --- | --- |
|  |  | Leaf 2 | Leaf 3 | Leaf 4 | Leaf 5 | Leaf 6 | Leaf 7 | Leaf 8 |
| *Oryza sativa ssp. japonica* | Nipponbare | 3.94 ± 0.35 (a) | 11.80 ± 1.15 (a) | 21.96 ± 1.53 (a) | 26.92 ± 2.00 (a) | 38.11 ± 1.73 (a) | 44.37 ± 1.40 (a) | 48.71 ± 2.84 (a) |
| *Oryza sativa ssp. indica* | IR 64 | 5.60 ± 0.22(b) | 11.75 ± 0.85 (a) | 20.95 ± 1.91 (a) | 26.25 ± 1.72 (a) | 34.05 ± 1.07 (a) | 41.56 ± 1.48 (b) | 44.85 ± 2.75 (b) |
| *Oryza sativa ssp. indica* | Vandana | 4.54 ± 0.62 (a) | 11.50 ± 1.43 (a) | 21.12 ± 1.55 (a) | 29.06 ± 1.34 (a) | 32.59 ± 1.71 (b) | 37.36 ± 1.77 (c) | 45.59 ± 1.74 (ab) |
| *Oryza sativa ssp. indica* | Swarna | 4.70 ± 0.44 (a) | 14.93 ± 1.24 (b) | 27.92 ± 2.52 (b) | 39.88 ± 2.65 (b) | 46.90 ± 1.87 (c) | 52.76 ± 2.86 (d) | 58.72 ± 3.03 (c) |
| *Oryza sativa ssp. indica* | Pusa Basmati 1 | 6.25 ± 0.54(b) | 14.66 ± 2.36 (b) | 28.48 ± 3.11 (b) | 37.11 ± 2.01 (c) | 42.98 ± 1.76 (d) | 48.30 ± 2.26 (a) | 55.17 ± 1.94 (d) |
| *Oryza sativa ssp. indica* | Nagina 22 | 4.21 ± 0.38 (a) | 14.55 ± 1.52 (b) | 27.98 ± 1.30 (b) | 38.50 ± 2.29 (bc) | 44.56 ± 2.14 (de) | 53.41 ± 2.59 (d) | 58.77 ± 1.79 (c) |
| *Oryza glaberrima* | IR 102925 | 6.00 ± 0.30(b) | 14.95 ± 1.65 (bc) | 26.90 ± 1.18 (b) | 38.20 ± 1.19 (c) | 45.90 ± 2.42 (ce) | 53.70 ± 2.40 (d) | 59.50 ± 2.82 (c) |
| *Oryza glumaepatula* | IR104151 | 8.01 ± 0.56 (c) | 23.24 ± 1.68 (e) | 37.26 ± 3.10 (ce) | 43.30 ± 1.30 (d) | 46.76 ± 1.51 (ce) | 53.35 ± 2.05 (d) | 63.91 ± 3.07 (e) |
| *Oryza punctata* | IRGC105137 | 6.08 ± 0.63(b) | 17.67 ± 1.19 (cd) | 34.38 ± 1.27 (cd) | 47.52 ± 1.39 (e) | 51.92 ± 1.31 (f) | 58.26 ± 1.23 (e) | 70.67 ± 2.04 (f) |
| *Oryza latifolia* | IRGC 99596 | 10.15 ± 0.62(d) | 18.32 ± 1.42 (d) | 31.52 ± 2.05 (d) | 46.13 ± 1.19 (e) | 53.12 ± 1.10 (f) | 60.63 ± 2.26 (e) | 70.93 ± 1.68 (f) |
| *Oryza rufipogon* | IRGC 99562 | 8.57 ± 1.57(c) | 23.75 ± 3.20 (e) | 37.55 ± 2.80 (e) | 47.00 ± 4.47 (e) | 66.40 ± 6.29 (g) | 74.72 ± 3.55 (f) | 85.56 ± 1.67 (g) |
| *Oryza australiensis* | IRGC 105272 | 20.57 ± 2.68 (e) | 34.19 ± 3.44 (f) | 50.66 ± 4.55 (f) | 56.57 ± 2.60 (f) | 64.74 ± 4.36 (g) | 69.50 ± 0.71 (f) | 87.80 ± 5.27 (g) |

**Fig. 1.**
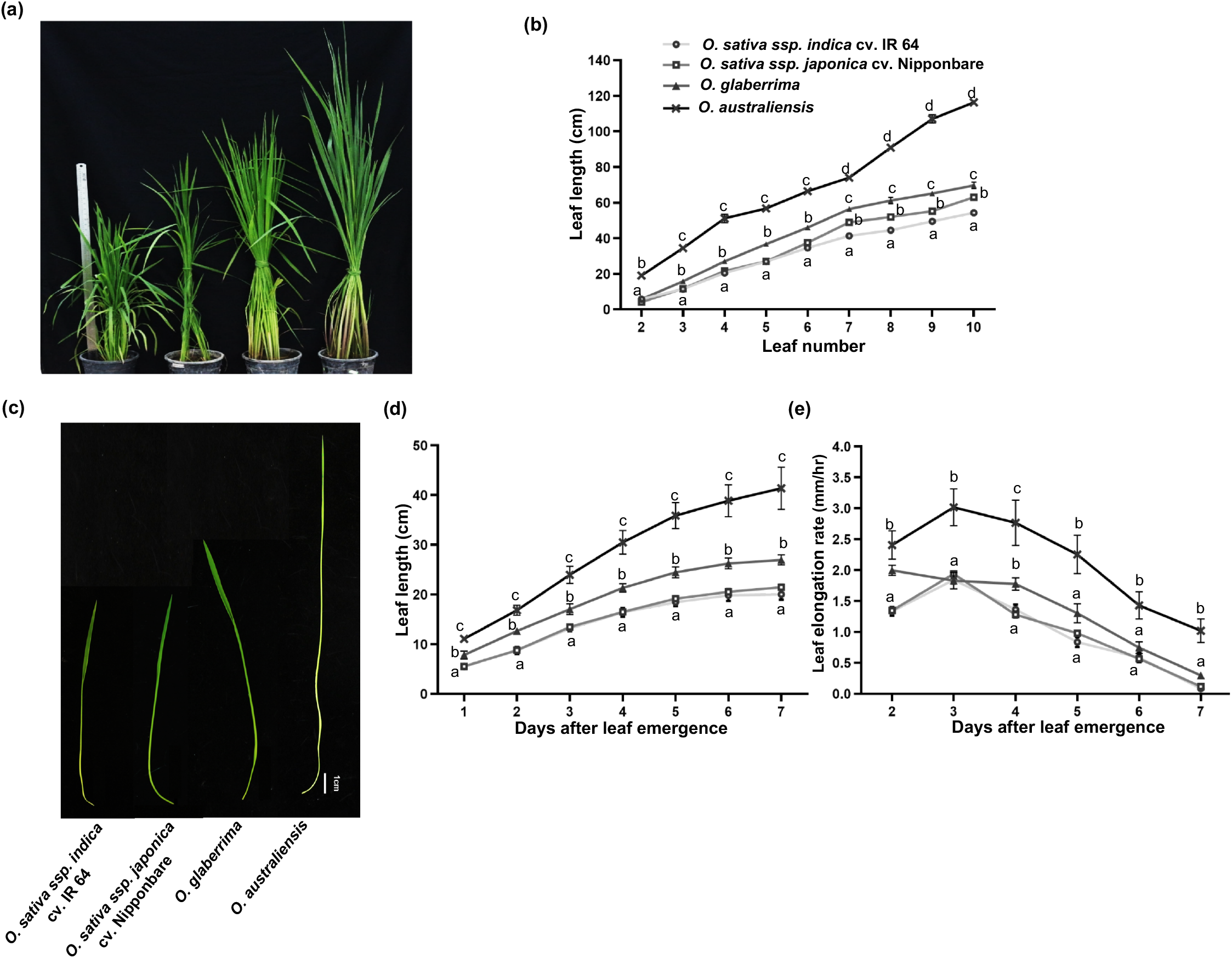
Wild rice *Oryza australiensis* grows longer leaf compared to the selected cultivated rice accessions. (a) Representative photographs of the selected cultivated and wild rice accessions showing leaf size variations. (b) Quantification of mature leaf length of the second leaf to the tenth leaf, (c) Representative fourth leaf images at the second day of emergence. (d) Quantification of fourth leaf length at the different days of emergence. (e) Leaf elongation rate of the selected rice accessions. Values in graphs represent mean±SD (n=15). Different letters indicate statistically significant differences according to one-way ANOVA followed by post-hoc Tukey HSD at P < 0.05.

Quantification of the fourth leaf length of the selected four accessions consistently showed longer leaf in the wild rice *O. australiensis* compared to the cultivated accessions at different days of emergence (Fig. 1c, d). *O. glaberrima* showed intermediate leaf length between the wild rice *O. australiensis* and cultivated Asian rice varieties at different days of emergence. Consistent with the leaf length, the leaf elongation rate (LER) was higher for the wild rice than the cultivated accessions at different days of emergence (Fig. 1e). Taken together, the wild rice *O. australiensis* showed remarkably longer leaves than the cultivated varieties, and *O. glaberrima* also developed significantly longer leaves than IR64 and Nipponbare.

### Transcriptome profiling identified cell cycle, GA, and GRFs as the key regulators of rice leaf length

To identify genes and pathways controlling the leaf length differences, we compared the transcriptomes of the developing fourth leaves of the selected accessions. The maximum number of differentially expressed genes (DEGs) was found between Nipponbare and *O. australiensis*, whereas a lower number of DEGs was found between cultivated accessions (Table S3, Dataset S1). Since the leaves of the wild rice *O. australiensis* were remarkably longer than IR64 and Nipponbare, we first compared the gene expression profiles between the leaves of *O. australiensis* and two cultivated varieties of *O. sativa*. 2,384 genes were expressed at higher levels and 3,870 genes were expressed at lower levels in growing leaves of *O. australiensis* compared to both IR64 and Nipponbare (Fig. 2a). The genes expressed at higher levels showed enrichment of GO terms ‘cell cycle’, ‘development process’, ‘cell growth’, and ‘cell morphogenesis’ (Dataset S2). The enriched GO terms for the genes expressed at lower levels included ‘photosynthesis’, ‘biosynthetic process’, and ‘signaling’. We also compared the gene expression differences between the significantly longer leaves of *O. glaberrima* and the two cultivated varieties of *O. sativa*. The 1,125 genes expressed at higher levels showed enrichment of GO terms ‘cell cycle’, ‘development process’, ‘growth’, and ‘hormone transport’, whereas 1,290 genes expressed at lower levels were enriched for the term ‘metabolic process’ (Fig. S2a; Dataset S2). We then analyzed the differentially expressed genes in the wild rice *O. australiensis* compared to all three cultivated accessions. The 1,707 shared genes expressed at higher levels in *O. australiensis* compared to all three cultivated accessions showed enrichment of GO terms “development process”, “cell cycle”, and “growth”, whereas the GO term “biosynthetic process” was enriched for genes expressed at lower levels (Fig. S2b; Dataset S2).

**Fig. 2.**
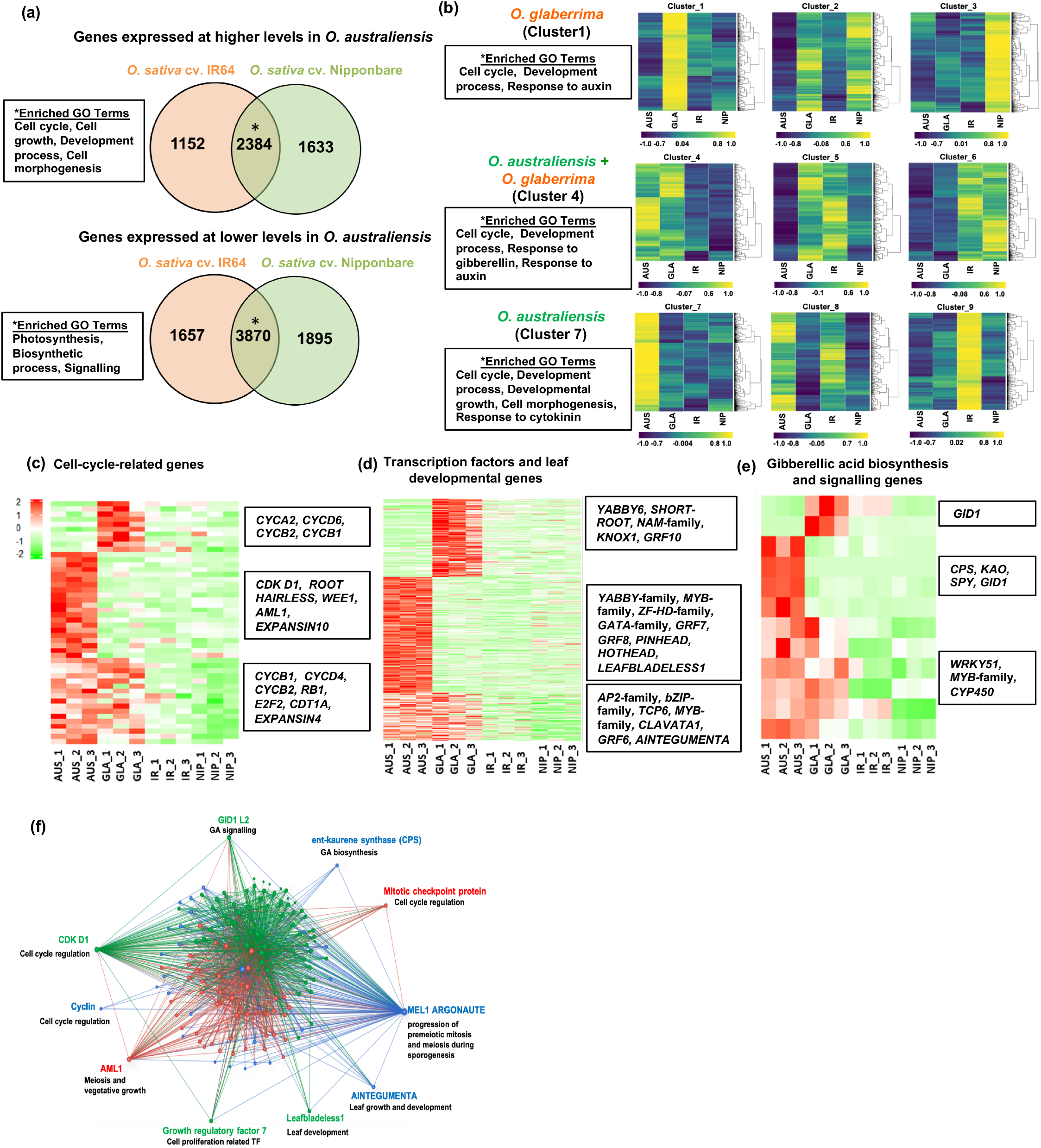
Transcriptome profiling reveals the key contribution of the cell cycle and Gibberellic Acid (GA) for differential leaf length across the selected rice accessions. (a) The number of differentially expressed transcripts and enriched GO terms in *O. australiensis* compared to *O. sativa* cv. IR64 and *O. sativa* cv. Nipponbare. * show enriched GO terms. (b) Gene expression clusters showing genotype-specific resolution for the selected four accessions derived from PCA-SOM analysis. Enriched GO terms are shown for *O. australiensis* and *O. glaberrima* specific clusters 1, 4, and 7. (c-e) Expression patterns of cell cycle-related genes (c), transcription factors and leaf developmental genes (d), and GA biosynthesis and signaling genes (e) present in clusters 1, 4 and 7 across the four accessions. (f) Gene regulatory network generated for the genes from the relevant clusters of (b). Three different modules of the network are shown in different colors. AUS= *O. australiensis*, GLA= *O. glaberrima*, IR*= O. sativa* cv. IR64, NIP= *O. sativa* cv. Nipponbare.

Principle component 1 (PC1), explaining 39% variation in the dataset, primarily explained the expression differences of the wild rice *O. australiensis* compared to the three cultivated accessions (Fig. S3a, b). PC2, explaining 31% of the overall variation, explained the expression differences of *O. glaberrima* compared to the other three accessions. The clusters generated by PCA-SOM provided the accession-specific resolution of the transcripts (Fig. 2b; Dataset S3). Since *O. australiensis* and *O. glaberrima* had significantly longer leaves compared to IR64 and Nipponbare, we analyzed the genes and enriched pathways in the clusters specific to either *O. australiensis* or *O. glaberrima* or both. The cluster 4 that had genes specific to both *O. australiensis* and *O. glaberrima* showed enrichment of terms ‘cell cycle’, ‘development process’, ‘response to gibberellin’, and ‘response to auxin’ (Fig. 2b; Dataset S4). The transcripts in *O. australiensis* specific cluster 7 and *O. glaberrima* specific cluster 1 were enriched for ‘development process’ and ‘cell cycle’.

Since transcription factors, phytohormones, and cell division and expansion are crucial for leaf size regulation, we compared the expression of the related genes from the relevant clusters across the selected accessions (Dataset S5). Several cell cycle genes, such as *CYCLINB*, *CYCLINA*, and *CDKD1*, as well as cell cycle regulators, such as *E2F2* and *RB1*, were expressed at higher levels in *O. australiensis* and *O. glaberrima* than IR64 and Nipponbare (Fig. 2c). Further, transcription factor genes known to regulate leaf growth and development, such as *GRF*-family, *TCP*-family, *YABBY*-family, *NAM-family*, and *ANT*, were expressed at higher levels in *O. australiensis* and *O. glaberrima* (Fig. 2d). Among the *GRFs*, *OsGRF7* and *OsGRF8* specifically showed higher expression in *O. australiensis* leaves compared to the cultivated varieties. GA biosynthesis and signaling genes that included *CPS, KAO*, and *GID1* were expressed at higher levels in *O. australiensis* and *O. glaberrima* than IR64 and Nipponbare (Fig. 2e). Moreover, differential expressions of auxin and cytokinin biosynthesis and signaling genes were also observed across the selected accessions (Fig. S3c).

The coexpression network of genes from the relevant clusters showed 3 major modules including more than 20 nodes (Fig. 2f; Dataset S6). The network showed significant connections for GA biosynthesis and signaling genes *CPS* and *GID1L2*, cell cycle-related genes *CYCLIN* and *CDKD1*, as well as *GRF7*. Investigation of the hub genes in module 2 showed *CDKD1* as one of the highly connected genes, which was connected to GA receptor *GID1L2*, and *GRF7* (Fig. S3d). Taken together, transcript profiling along with gene co-expression network highlighted cell cycle, GA, and transcription factors, such as OsGRF7 and OsGRF8, as key contributors of rice leaf length regulation.

### GA-levels correlated with the leaf length variation across the cultivated and wild rice accessions

Since the transcript profiling indicated the involvement of GA for the leaf length differences across the accessions, we quantified GA in the fourth leaves of the cultivated and wild rice accessions that showed significant variation for leaf length (Fig. S4). We detected significantly higher levels of GA_4_ in the leaves of wild rice accessions with longer leaves compared to the cultivated rice accessions (Fig. 3a). A strong significant positive correlation between leaf length and GA_4_ levels was observed for the selected cultivated and wild rice accessions, suggesting a key role of GA in rice leaf length regulation (Fig. 3b). We further detected a higher level of GA_7_ in addition to GA_4_ in the leaves of *O. australiensis* (wild rice with longest leaves among all the investigated accessions) compared to Nipponbare (cultivated rice variety with smaller leaves) (Fig. 3c).

**Fig. 3.**
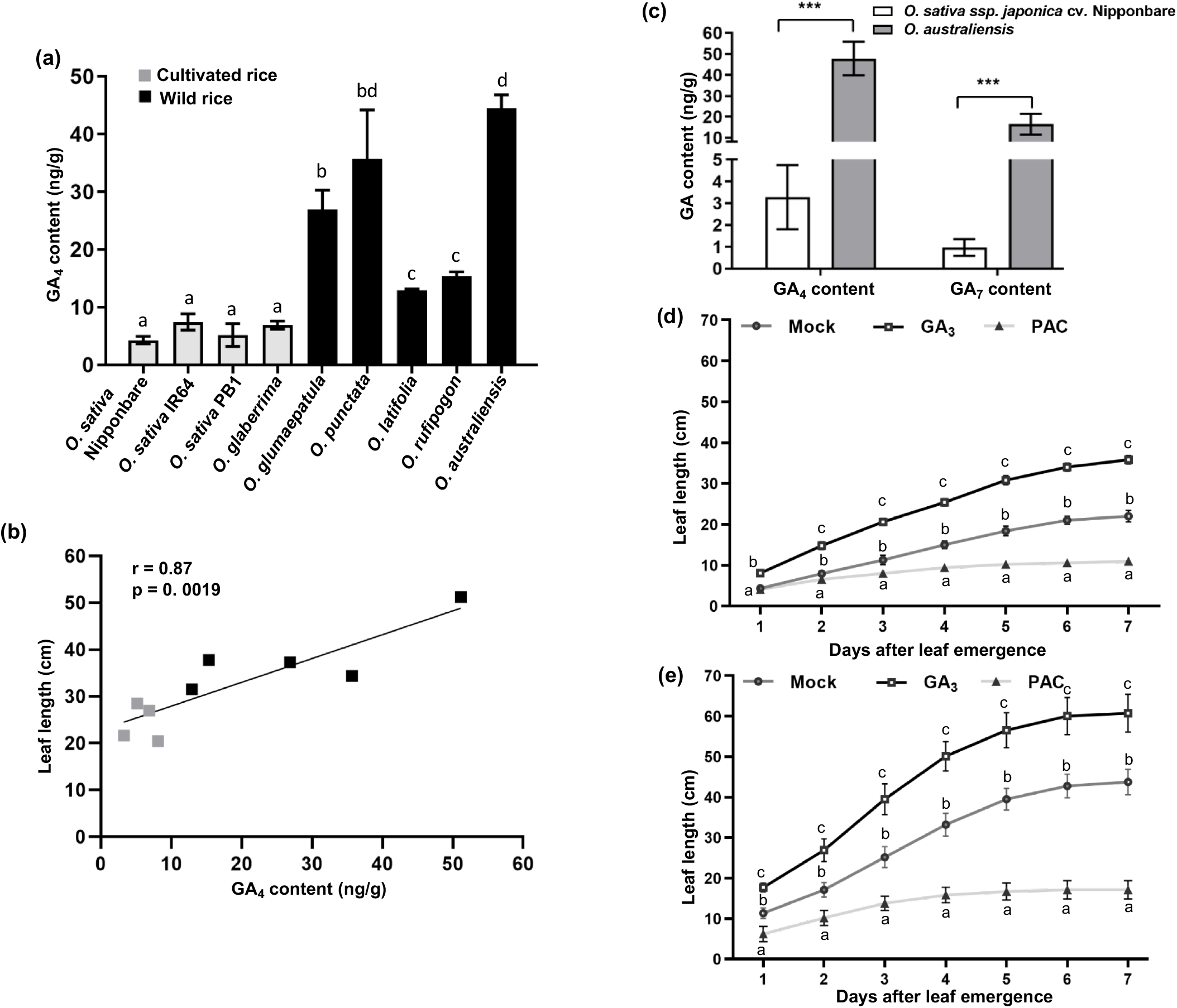
GA levels correlate with the leaf length variation across the cultivated and wild rice accessions. (a) GA_4_ quantification in the fourth leaf of the cultivated and wild rice accessions on the second day of emergence. Data show mean +SD (n=5). (b) Correlation between GA_4_ and leaf length. (c) GA_4_ and GA_7_ quantification in the fourth leaf of *O. sativa* cv. Nipponbare and *O. australiensis* on the second day of emergence. Data show mean +SD (n=5),*** P□<□ 0.001 by Student’s Etest. (d, e) Effects of GA_3_ (10μM) and PAC (1μM) treatments on leaf length of *O. sativa* cv. Nipponbare (d) and *O. australiensis* (e). Data show mean +SD (n=15). Different letters indicate statistically significant differences according to one-way ANOVA followed by post-hoc Tukey HSD at P < 0.05.

To confirm the GA-mediated rice leaf length regulation, we examined the effects of exogenous treatment of GA_3_ and GA-biosynthesis inhibitor Paclobutrazol (PAC) on leaf length of the two contrasting accessions, *O. australiensis* and Nipponbare. GA_3_ treatment promoted the leaf length of both Nipponbare and *O. australiensis* (Fig. 3d, e, and S5a, b). The extent of increase in leaf length was more in Nipponbare (~1.6-fold) than *O. australiensis* (~1.3-fold). Exogenous treatment of PAC drastically reduced the leaf length of both the accessions. PAC inhibited the leaf length of Nipponbare by ~2-fold, and of *O. australiensis* by ~2.5-fold. The higher extent of leaf length increase in Nipponbare by GA_3_ treatment and higher extent of leaf length decrease in *O. australiensis* under PAC along with a general higher GA content in wild rice accessions with longer leaves suggested that GA levels could be determining for the differences in the leaf length across the cultivated and wild rice accessions.

### Leaf kinematics confirmed the GA-mediated control of division zone to regulate rice leaf length

Since transcriptomic comparison showed higher expression of cell-cycle related genes in the wild rice accessions than the cultivated varieties, we performed DAPI staining and kinematics analyses on the growing fourth leaves to investigate the differences in cell division and growth zones across the selected accessions. *O. australiensis*, with longer leaves than the cultivated varieties, showed dividing cells for longer length from the leaf base, and thus longer division zone at different days of emergence (Fig. S6a). *O. glaberrima* also showed a longer division zone compared to IR64 and Nipponbare. Detailed kinematics analyses on the fourth leaves of each selected accession confirmed the differences in the division zone (Fig. 4a, b, and S6; Table 2). All the four selected accessions showed a sigmoidal curve for epidermal cell length profile, consisting of a division zone at leaf base followed by elongation zone and maturation zone (Fig. 4a, b and S6f, g). The cultivated varieties Nipponbare and IR64 showed division zone extending up to 7.8mm and 7.5mm from the leaf base, respectively (Fig. 4a and S5b, d, f). *O. australiensis*, with the longest leaves, showed a remarkably longer division zone that extended to 16.7mm from the leaf base (Fig. 4b and S5c, e). The longer division zone in *O. australiensis* leaves was accompanied by a faster cell production rate and leaf elongation rate (Table 2). *O. glaberrima* also showed a longer division zone as well as a higher cell production rate compared to IR64 and Nipponbare (Fig. S5g).

**Fig. 4.**
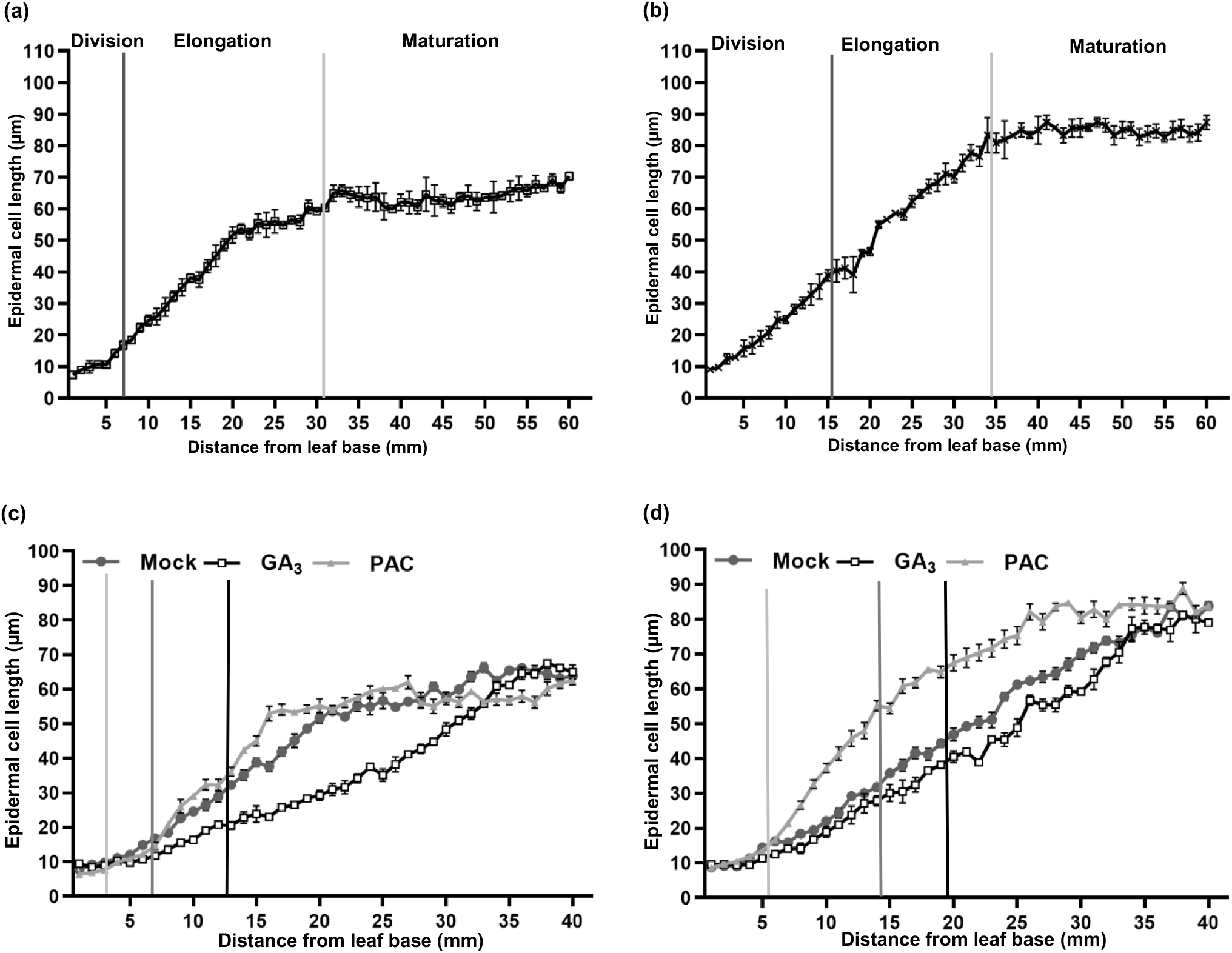
Leaf kinematics analyses confirm the GA-mediated control of cell division for rice leaf length regulation. (a, b) Different growth zones on the fourth leaf of *O. sativa* cv. Nipponbare (a) and *O. australiensis* (b) at the second day of emergence as quantified by epidermal cell length profiling and DAPI staining. Vertical lines of different shades represent the separation of different leaf-growth zones. (c, d) Different growth zones on the fourth leaf of *O. sativa* cv. Nipponbare (c) and *O. australiensis* (d) in response to GA_3_ and PAC treatments at the second day of emergence as quantified by epidermal cell length profiling and DAPI staining. Vertical lines of different shades represent the size of division zones under different treatments. Values in graphs represent mean±SD (n=5).

**Table 2.** Quantification of leaf kinematics parameters of the growing fourth leaf of the selected cultivated and wild rice accessions. Each value represents mean ± SD (n = 15 for mature leaf length and leaf elongation rate, n = 5 for all other parameters). Different letters indicate statistically significant differences according to one-way ANOVA followed by post-hoc Tukey HSD calculation at P < 0.05.

| Growth parameters | *O. sativa* cv. IR 64 | *O. sativa* cv. Nipponbare | *O. glaberrima* | *O. australiensis* |
| --- | --- | --- | --- | --- |
| Mature leaf length (cm) | 20.06 ± 1.38(a) | 21.46 ± 0.81(a) | 26.98 ± 1.01(b) | 41.38 ± 4.25(c) |
| Leaf elongation rate (mm/h) | 1.32 ± 0.01(a) | 1.34 ± 0.01(a) | 2.01 ± 0.01(b) | 2.41 ± 0.01(b) |
| Meristem length (mm) | 7.52 ± 0.14(a) | 7.81 ± 0.34(a) | 12.32 ± 0.34(b) | 16.66 ± 0.64(c) |
| Length of the growth zone (mm) | 24.00 ± 1.00(a) | 31.33 ± 0.58(b) | 24.67 ± 1.53(a) | 34.33 ± 0.58(c) |
| Mature cell length (μm) | 70.61 ± 0.15(a) | 64.26 ± 1.41(b) | 66.00 ± 1.07(b) | 84.57 ± 1.32(c) |
| Cell production rate (cells/h) | 18.76 ± 0.12(a) | 20.88 ± 0.40(b) | 30.44 ± 0.51(c) | 28.49 ± 0.42(d) |
| Number of cells in the meristem | 738.29 ± 27.50(a) | 715.92 ± 17.05(a) | 842.73 ± 31.50(b) | 904.75 ± 52.90(b) |
| Number of cells in the growth zone | 1193.63 ± 70.37(a) | 1328.80 ± 26.33(b) | 1100.53 ± 43.44(ac) | 1211.09 ± 38.48(abc) |
| Number of cells in the elongation zone | 455.35 ± 68.50(a) | 612.88 ± 41.50(b) | 257.80 ± 38.30(c) | 306.34 ± 19.50(c) |
| Average cell division rate (cell. cell<sup>-1</sup>.h<sup>-1</sup>) | 0.03 ± 0.001(a) | 0.03 ± 0.001(ab) | 0.04 ± 0.001(bc) | 0.03 ± 0.001(ab) |
| Cell cycle duration (h) | 27.29 ± 1.15(a) | 23.78 ± 0.99(b) | 19.20 ± 1.03(c) | 22.00 ± 0.98(b) |
| Time in the elongation zone (h) | 24.26 ± 3.53(a) | 29.34 ± 1.62(a) | 8.47 ± 1.29(b) | 10.76 ± 0.79(b) |
| Time in division zone (h) | 260.01 ± 12.40(a) | 225.54 ± 10.10(b) | 186.66 ± 11.05(c) | 216.13 ± 11.45(bc) |
| Length of cells leaving meristem (μm) | 15.15 ± 0.37(a) | 17.79 ± 0.41(a) | 31.33 ± 1.41(b) | 39.93 ± 3.36(c) |
| Average cell expansion rates (μm. μm<sup>-1</sup>.h<sup>-1</sup>) | 0.06 ± 0.01(ac) | 0.04 ± 0.003(a) | 0.09 ± 0.01(bc) | 0.07 ± 0.01(ac) |

We then asked the question of whether GA regulates rice leaf length by controlling cell proliferation and size of the division zone. To this end, we performed leaf kinematics analysis on the fourth leaves of *O. australiensis* and Nipponbare under GA_3_ and PAC treatment. We observed that the leaf length promotion in response to GA treatment was due to an increase in the size of the division zone in both the accessions (Fig. 4c, d, and S5c, d). However, the impact of GA on cell division was more striking in Nipponbare than *O. australiensis* in terms of the size of the division zone, cell production rate, and the number of cells in the meristem (Tables S4 and S5). The inhibitory effect of PAC on the leaf length was also clearly due to the reduced size of the division zone in both the accessions (Fig. 4c, d, and S5c, d). Interestingly, the kinematics profile of GA-treated Nipponbare leaves resembled that of *O. australiensis* leaves under control conditions (Fig. 4b, c). Together, the leaf kinematics profile of the contrasting rice accessions and the effects of exogenous GA and PAC treatments on the division zone confirmed that the higher GA levels promoted the rice leaf length by increasing the size of the division zone.

### Growth-zone-specific expression analysis of GA-related genes and associated GA levels

Next, we asked if zone-specific expressions of GA-biosynthesis and signaling genes along with associated GA levels explain the leaf size differences. To this end, we quantified the expression of GA biosynthesis genes in two leaf zones of Nipponbare and *O. australiensis:* zone I (leaf base – 0.8 cm; division zone in both the accessions) and zone II (0.8 – 1.6cm; elongation zone in Nipponbare, where division extends in *O. australiensis)*. The transcript levels of GA biosynthetic genes *OsGA20OX2* and *OsGA3OX2* were higher in the zone I of *O. australiensis*, which showed longer leaves and had higher GA levels in their leaves than Nipponbare (Fig. 5a, b). Furthermore, the expression level of the GA catabolic gene *OsGA2OX4* was lower in zone II of *O. australiensis* than Nipponbare (Fig. 5c). Consistent with the expression of GA-biosynthesis and catabolic genes, more GA_4_ and GA_7_ levels were detected in both zones of *O. australiensis* than Nipponbare (Fig. 5e, f). The transcript levels of *OsSLR1*, encoding a GA-signalling repressor DELLA protein, was also higher in both the regions of Nipponbare leaves (Fig. 5d). Taken together, higher expression of GA biosynthetic genes *OsGA20OX2* and *OsGA3OX2* in zone I, lower expression of GA catabolic gene *OsGA2OX4* in zone II, and lower expression of GA-signalling repressor *OsSLR1* in *O. australiensis* promoted more zone-specific GA levels, and thus higher GA effects in *O. australiensis* than Nipponabre.

**Fig. 5.**
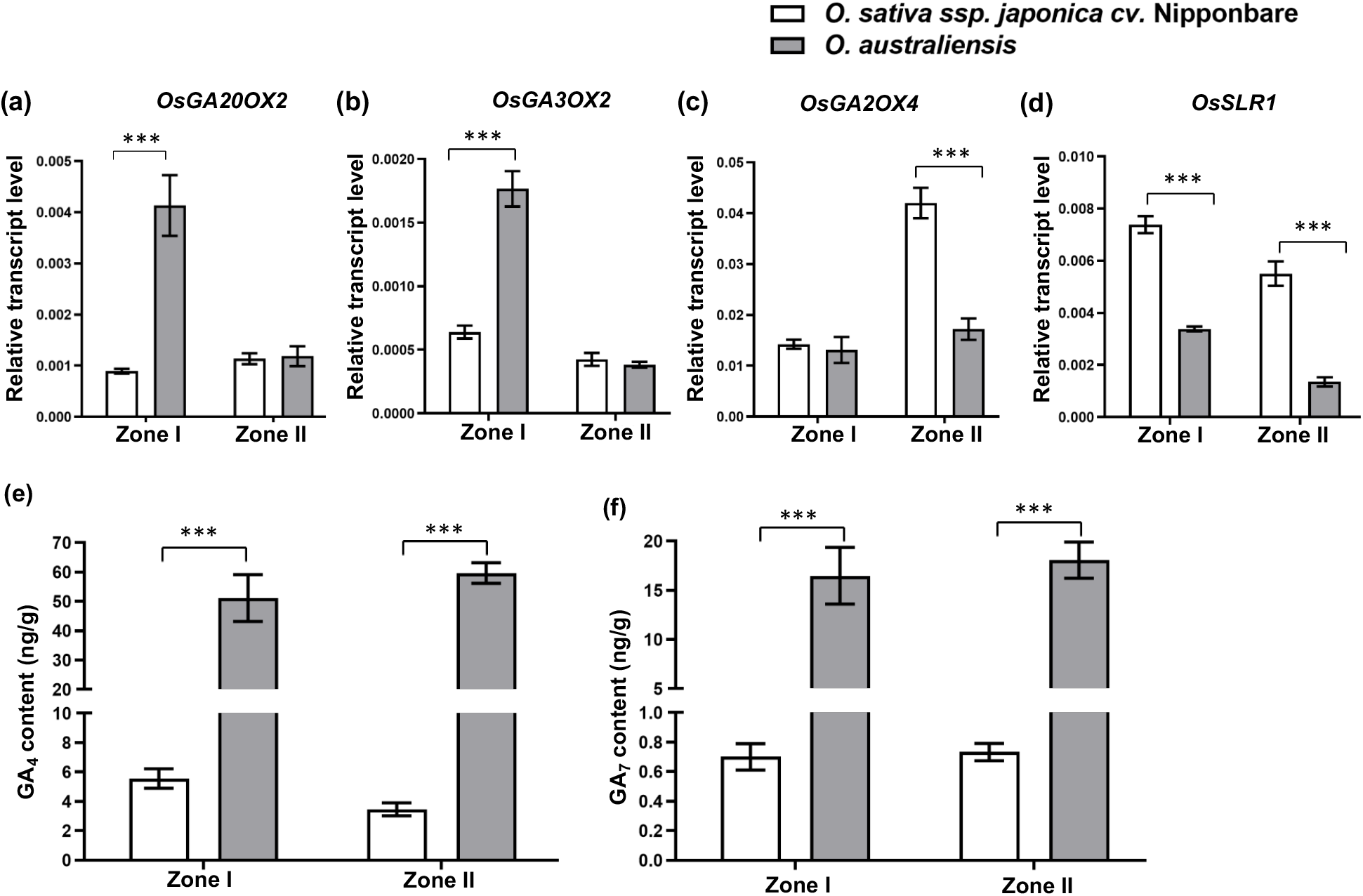
Zone-specific expression analysis of GA-related genes and GA quantification. (a - d) Expression pattern of GA biosynthesis genes *OsGA20OX2* and *OsGA3OX2* (a, b), GA catabolic gene *OsGA2OX4* (c), and GA-signalling repressor *OsSLR1* (d) in the two growth zones of fourth leaves of *O. sativa* cv. Nipponbare and *O. australiensis* on the second day of emergence. Data shown are mean +SD of three biological replicates, *** P□<□0.001 by Student’s t□test (e, f) Zone specific GA_4_ (e) and GA_7_ (f) quantification in the two growth zones of fourth leaves of *O. sativa* cv. Nipponbare and *O. australiensis* on the second day of emergence. Data shown are mean +SD (n=5),*** P□<□0.001 by Student’s t□test. Zone I represents the region from leaf base to 0.8cm and zone II represents 0.8cm to 1.6cm from the leaf base.

### Higher GA levels expanded the expression domain of *GRFs* and cell-cycle-related genes

Similar to the expression of GA-biosynthesis and signaling genes, we observed remarkably higher transcript levels of *OsGRF7* and *OsGRF8* in both zones I and II of *O. australiensis* leaves than Nipponbare (Fig. 6a, b). Further to corroborate the involvement of *OsGRF7* and *OsGRF8* in rice length regulation, we analyzed the transcripts level of the two *GRF*s in the two growth zones of cultivated and wild rice accessions initially used for leaf phenotyping and GA quantification. Wild rice accessions with longer leaves and higher GA content had a higher abundance of *OsGRF7* and *OsGRF8* transcripts in division zones (Fig. S7a, b). The transcript levels of *OsCYCB1;4* and *OsCYCA3;2* were also higher in both zone I and II of the *O. australiensis* than Nipponbare (Fig. 6c, d). Limited transcript levels of *OsCYCB1;4* and *OsCYCA3;2* in zone II of Nipponbare leaves matched with the negligible cell division activity beyond 0.8cm in the accession. The significantly lower expression level of cell cycle inhibitor *OsKRP4* and a higher level of cell cycle activator *OsE2F2* in basal regions of *O. australiensis* leaves than Nipponbare further reflected on the differences in cell cycle between the two accessions (Fig. S7c, d). The exogenous GA treatment remarkably induced the expression levels of *OsGRF7* and *OsGRF8* as well as *OsCYCB1;4* and *OsCYCA3;2* in zone II of Nipponbare (Fig. 6e-h). Moreover, the expression of *OsSLR1* was suppressed in both the zones in response to GA treatment (Fig. S8a). PAC treatment led to significant suppression of the expressions of *GRF*s and *CYCLIN*s in the division zone of Nipponbare (Fig. S8b). Together, gene expression analyses in different leaf zones of the selected cultivated and wild rice accessions as well as in response to the exogenous GA treatment strongly supported our hypothesis of GA-mediated cell cycle control through OsGRF7 and OsGRF8 to determine the rice leaf length.

**Fig. 6.**
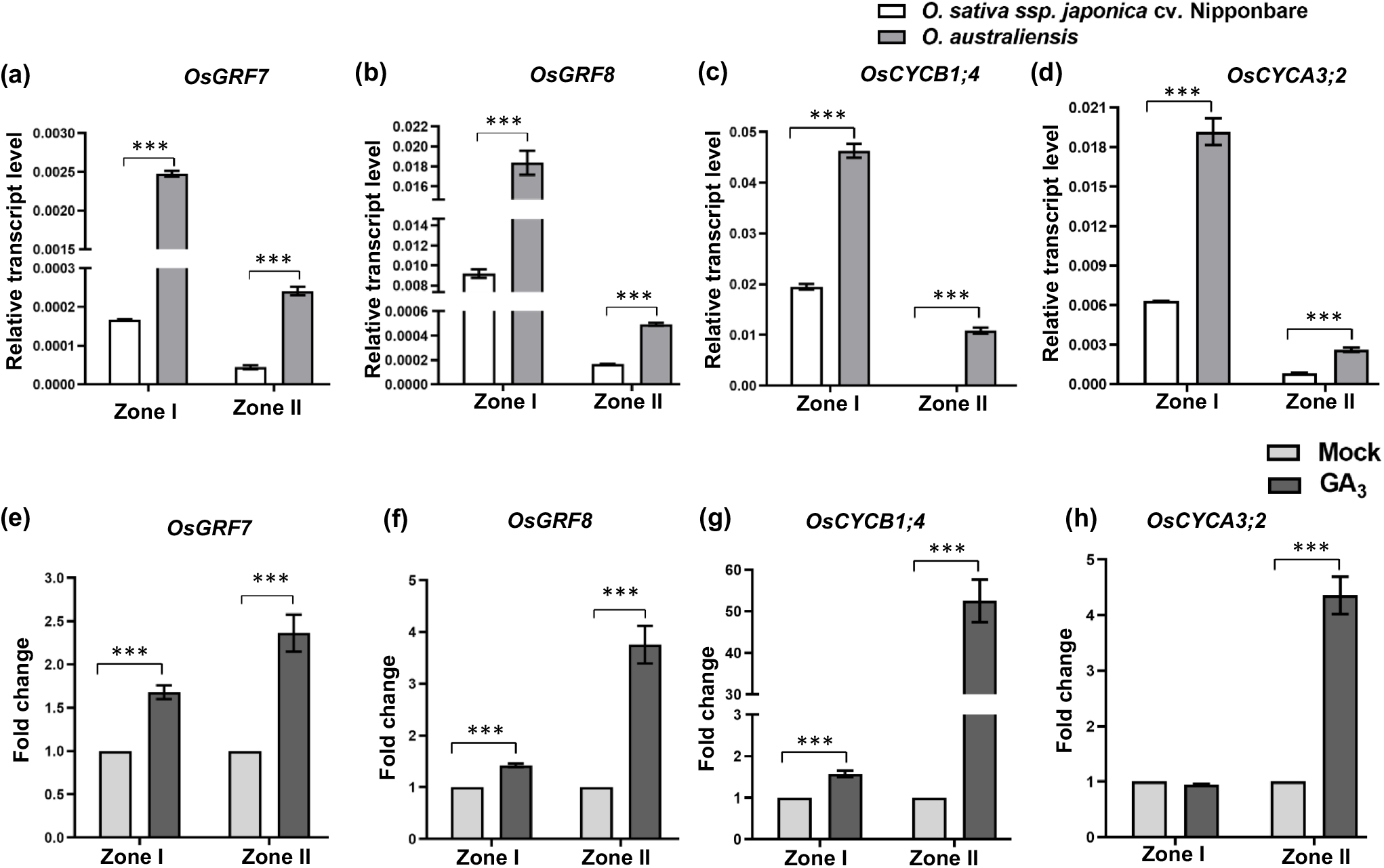
GA treatment affects zone-specific expression patterns of *GRFs* and cell cycle genes. (a - d) Expression pattern of genes encoding GRF transcription factors *OsGRF7* and *OsGRF8* (a, b), and cell cycle genes *OsCYCB1;4* and *OsCYCA3;2* (c, d) in the two growth zones of fourth leaves of *O. sativa* cv. Nipponbare and *O. australiensis* on the second day of emergence. (e - h) Effect of GA_3_ treatment on the expression of *OsGRF7* (e), *OsGRF8* (f)*, OsCYCB1;4* (g), and *OsCYCA3;2* (h) in the two growth zones of fourth leaves of *O. sativa* cv. Nipponbare. Data shown are mean +SD of three biological replicates, *** P□<□ 0.001 by Student’s t□test. Zone I represents the region from leaf base to 0.8cm and zone II represents 0.8cm to 1.6cm from the leaf base.

### Gene silencing confirmed the regulation of division zone size by GA and GRFs to control rice leaf length

To confirm the GA-mediated leaf length differences between the contrasting cultivated and wild rice accessions via specific GRFs, we performed transient silencing of genes encoding a GA biosynthesis enzyme GA20OX2, a GA signaling repressor SLR1, and Growth-Regulating Factors GRF7 and GRF8 in *O. australiensis* and Nipponbare. Since extended division zone and longer leaf length in the wild rice *O. australiensis* was associated with higher GA levels in leaves, we checked the effects of reducing GA levels by silencing *GA20OX2* in the wild rice. The silenced *O. australiensis* lines showed a remarkable reduction in the leaf length associated with decreased division zone size and lower expression of *OsGRF7*, *OsGRF8*, and *CYCLINs* compared to the control plants (Fig. 7a-c and S10a). Similar to *OsGA20OX2*, silencing of *OsGRF7* and *OsGRF8* in *O. australiensis* reduced the leaf length along with the smaller size of the division zone and reduced expression of downstream *CYCLIN* genes (Fig. 7a-c, and S10c). Silencing of *OsGA20OX2*, *OsGRF7*, and *OsGRF8* in Nipponbare also led to a reduction in leaf length, similar to exogenous PAC treatment, with reduced size of the division zone as well as lower expression levels of *OsGRF7, OsGRF8*, and *CYCLINs* (Fig. 7d-f and S10b, d). In contrast, silencing of *OsSLR1* in both the accessions led to a longer leaf, increased size of division zone, and higher expression levels of *OsGRF7*, *OsGRF8*, and *CYCLINs* compared to the control (Fig. S9e-h and S10e-f). Together, leaf phenotypes of *O. australiensis* and Nipponbare silencing lines confirmed the regulation of cell division zone size by GA and downstream OsGRF7 and OsGRF8 transcription factors to determine the rice leaf length.

**Fig. 7.**
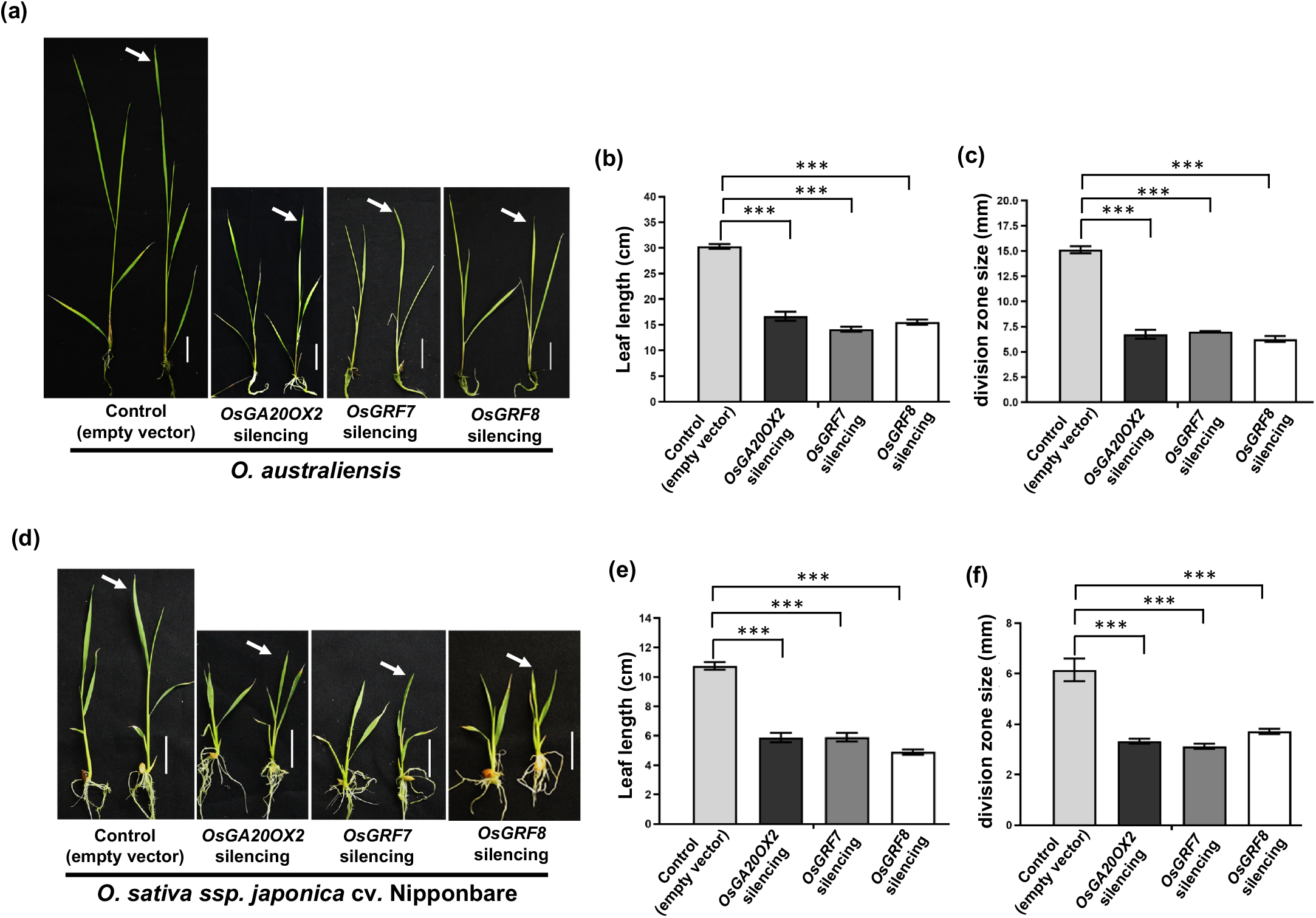
Silencing of GA-related genes confirms the GA-mediated regulation of division zone size to control rice leaf length. (a) Shown are the leaf phenotypes of *OsGA20OX2*, *OsGRF7*, and *OsGRF8* silencing in *O. australiensis* along with the control plants. (b, c) Effect of *OsGA20OX2, OsGRF7*, and *OsGRF8* silencing on leaf length (b) and size of division zone (c) in *O. australiensis*. (d) Shown are the leaf phenotypes of *OsGA20OX2*, *OsGRF7*, and *OsGRF8* silencing in *O. sativa* cv. Nipponbare along with the control plants. (e, f) Effect of *OsGA20OX2, OsGRF7*, and *OsGRF8* silencing on leaf length (e) and size of division zone (f) in *O. sativa* cv. Nipponbare. Data represent mean +SD of at least three biological replicates, *** P□<□0.001 by Student’s t-test. White arrow indicates fourth leaf and scale bar = 2cm in (a) and (d).

## Discussion

Natural variation for the leaf size in a crop species and related wild accessions is an excellent resource for understanding the genetic and hormonal basis of leaf size regulation. Studies have reported remarkable variations in the leaf morphological and anatomical features across cultivated and wild species of genus *Oryza*, suggesting a strong genetic control on leaf features of rice (Chatterjee *et al*., 2016; Mathan *et al*., 2021b). We performed detailed leaf phenotyping of seven cultivated and five wild rice accessions followed by GA quantification in leaves. Four accessions were then selected for detailed mechanistic studies that not only represented a significant genetic diversity but also covered the entire range of leaf length variations among the investigated accessions.

Differences in the leaf length are attributed to basic cellular features and leaf developmental programing (Gonzalez *et al*., 2012; Kierzkowski *et al*., 2019). Consistent with this, cell cycle and development were the obvious enriched terms for the genes expressed at higher levels in the accessions with longer leaves. Detailed leaf kinematics analyses for the contrasting cultivated and wild rice accessions precisely established the predominant role of cell division activity, and thus the size of the division zone, for defining the rice leaf length. Higher cell production rate and expansion of cell division to a longer length from leaf base drove the longer leaf phenotype in the wild rice *O. asutraliensis* than the cultivated varieties (Fig. 4; Table 2). Cell cycle duration and cell division rate were comparable across the accessions, suggesting that the longer cell division zone and increased cell production rate at the leaf base resulted in the production of a higher number of cells that contributed to longer leaves in the wild rice (Gazquez & Beemster, 2017). Higher expression levels and expanded expression domains of genes encoding cyclins and cyclin-dependent kinases in the growing leaves of *O. asutraliensis* than Nipponbare further supported the key importance of the cell division activity for the rice leaf length regulation (Fig. 2c and 6c, d). *O. asutraliensis*, besides showing the expansion of the division zone, showed longer epidermal cells compared to cultivated varieties, suggesting that the cell elongation also contributes to the longer leaf length of the wild rice (Fig. 4b).

Spatial and temporal regulation of cell division activity in plants is under complex regulation of phytohormones and transcription factors (Nelissen *et al*., 2012; Takatsuka & Umeda, 2014). We detected auxin-, cytokinin-, and GA-related genes among the differentially expressed genes for the selected rice accessions. Besides, genes encoding transcription factors known to regulate cell division, such as GRFs and TCPs, were also differentially expressed between the contrasting rice accessions (Vercruysse *et al*., 2020b). GA is shown to control the cell division activity via the involvement of GRF transcription factors in Arabidopsis and maize (Lantzouni *et al*., 2020; Fina *et al*., 2017). Our study shows that this mechanism could have been instrumental in optimizing rice leaf length during the domestication as wild rice species with longer leaves had higher endogenous GA levels than the cultivated varieties. GA- and GRF-mediated spatial control of cell division activity explain the difference in the size of the division zone leading to leaf length diversity in rice. Comparative transcriptome profiling on full leaves showed higher expression of GA-biosynthesis and signaling genes as well as GRF transcription factors in the longer leaves of *O. australiensis* and *O. glaberrima* than IR64 and Nipponbare, suggesting the key functions of GA and downstream GRFs to regulate the division zone size (Fig. 2d, e). Given the spatial genetic regulation for the leaf size control, functions of these factors at the leaf basal regions with cell division activity could be critical to the final leaf size. Consistent with this, our results on zone-specific expression profiling supported the key role of local differences in the expression of the relevant genes at the leaf basal region for leaf size regulation.

In support of the GA-mediated regulation of rice leaf length by expanding cell division activity, growing leaves of *O. australeinsis* had higher levels of endogenous GAs in the basal division zone than Nipponbare (Fig. 5 e, f). Higher expression levels of GA biosynthesis genes *OsGA2OOX2* and *OsGA20OX2* in the basal division zone of *O. australeinsis* leaves than Nipponbare corroborated the higher GA content in the wild rice (Fig. 5a, b). In addition, lower expression of GA-catabolic gene *OsGA2OX4* in the transition zone of the wild rice than Nipponbare likely maintained higher GA levels in the zone. Consistent with these, exogenous GA-treatment increased the rice leaf length by extending the zone where cell division operates. GA is known to induce cell elongation response (Jones & Kaufman, 1983). Exogenous GA-treatment, however, increased the leaf length without altering the mature epidermal cell length, suggesting that GA primarily affects rice leaf length via regulation of cell division (Tables S4 and S5). Finally, reduction in leaf length of both cultivated rice Nipponbare and wild rice *O. australiensis* by silencing *OsGA20OX2* that involved reduced division zone and downregulation of *OsGRF7*, *OsGRF8*, and *CYCLIN*s confirmed the central role of GA in rice leaf length regulation through the control of the size of division zone (Fig. 7a-f and S10a, b).

GRF transcription factors promote cell division activity by directly binding to the promoters of *CYCA1;1* and *CDC20* (Li *et al*., 2018). OsGRF7 regulation of GA-biosynthesis and auxin-signaling genes control the rice plant architecture (Chen *et al*., 2020). In addition, OsGRF1 is shown to influence rice leaf growth (van der Knaap *et al*., 2000). We identified specific OsGRFs regulating cell division activity to control the size of the division zone for final rice leaf size determination by genetic comparison of wild rice with longer leaves with the cultivated varieties. Higher *OsGRF7* and *OsGRF8* expression in growing leaves of *O. australiensis* with longer leaves from RNAseq analysis along with more *OsGRF7* and *OsGRF8* transcripts in division zone of wild rice accessions than cultivated varieties suggested that *Os*GRF7 and *Os*GRF8 may connect GA-levels and size of division zone for rice leaf length regulation (Fig. 6a, b and S7a, b). Interestingly, *OsGRF7* and *OsGRF8* were found to be closely related to *AtGRF1* and *AtGRF2*, which are shown to influence Arabidopsis leaf size, in phylogenetic analysis (Kim *et al*., 2003; Zhang et al., 2008; Chen *et al*., 2020). GA-treatment, increasing the leaf length by increasing the domain of cell division, induced the *OsGRF7* and *OsGRF8* expression in the division zone, further, suggesting the OsGRF7 and OsGRF8 functions downstream to GA (Fig. 6e-h). Consistent with this, silencing of *OsGRF7* and *OsGRF8* resulted in reduced rice leaf length with an associated decrease in the expression of the *CYCLINs* (Fig. 7 and S10c, d). Silencing of *OsSLR1*, a DELLA protein that inhibits GRF functions to repress GA signaling and whose expression is suppressed by exogenous GA treatment, led to increased leaf length along with higher expression of *OsGRF7* and *OsGRF8* further supporting the functions of OsGRF7 and OsGRF8 to integrate GA levels and size of division zone to regulate rice leaf length (Fig. S9e-h).

In conclusion, we present a detailed cellular basis and underlying genetic mechanism for leaf length regulation in rice, a staple diet for more than half of the world’s population. We showed that the size of the cell division zone regulated by GA and downstream OsGRF7 and OsGRF8 transcription factors determines the final rice leaf length and explains the variation in leaf length across the cultivated and wild rice accessions (Fig. 8). Higher GA levels and increased abundance of *OsGRF7* and *OsGRF8* resulting in expanded cell division zone explained the longer leaves of the wild rice accessions compared to the cultivated varieties (Fig. 8). The established GA-OsGRF7/OsGRF8 module to control the division zone size could not only be a general regulator of rice leaf length but might also have contributed to the optimization of leaf size during domestication. The module could, further, be a way for plants to achieve cellular plasticity in response to their growth environment.

**Fig. 8.**
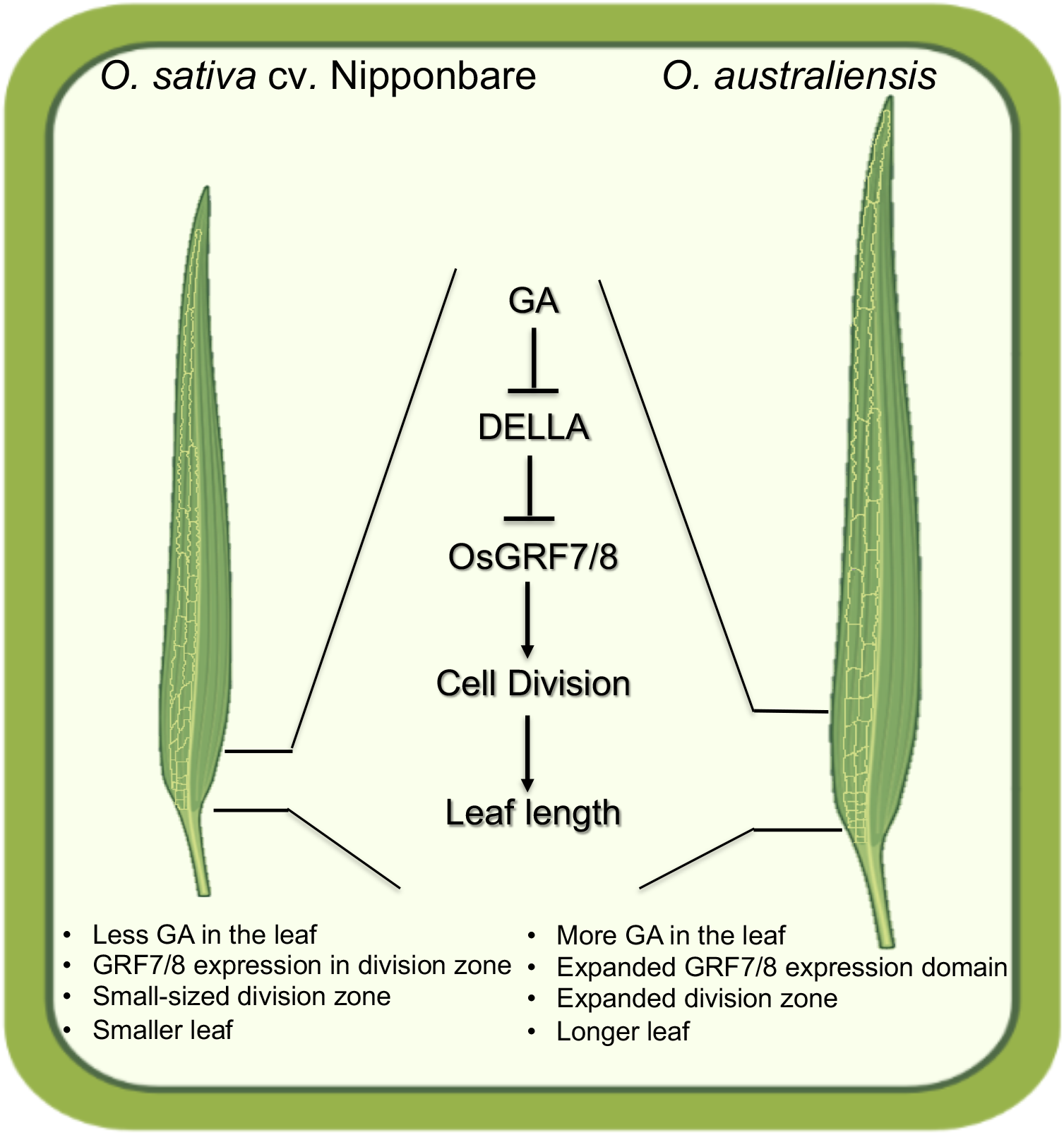
GA-OsGRF7/8 module is a key regulator of the rice leaf length via controlling the size of the division zone. GA-mediated induction in the expression domain of OsGRF7 and OsGRF8 transcription factors expands the region with cell division activity at the leaf base, thus promoting the longer division zone with a higher cell production rate leading to longer leaves in rice. Higher GA levels and downstream signaling resulting in increased cell division activity explained the longer leaves of the wild rice *O. australiensis* compared to the cultivated varieties.

## Acknowledgments

This work was supported by the core funding from the National Institute of Plant Genome Research as well as Ramalingaswamy Re-entry Fellowship (BT/RLF/Re-entry/05/2013) and Rice Network Project (BT/Ag/Network/Rice/2019-20) from the Department of Biotechnology, Ministry of Science and Technology, India. V.J. and K.S. acknowledge their UGC-SRF and SERB-NPDF fellowships, respectively. We thank the NIPGR Metabolomics Facility for Gibberellic Acid (GA) quantification and NIPGR Central Instrumentation Facility for their support. Seeds of wild rice species and *Oryza glaberrima* were kindly provided by Dr. Kuldeep Singh and Dr. Kumari Neelam, Punjab Agricultural University, Ludhiana, India.

## Author contributions

V.J. and A.R, conceptualized the study. V.J., K.S., and A.R. designed experiments. V.J. and A.C. performed experiments. V.J., R.R., Y.I., and A.R. analyzed transcriptomic data. V.J. and K.S. analyzed leaf kinematics data. V.J. and A.R. compiled figures and wrote the manuscript. V.J., K.S. and A.R. edited the final manuscript with the contribution of all the authors.

## Data Availability Statement

The data that supports the findings of this study are available in the supplementary material of this article. The quality-filtered reads, which were used to get the normalized read counts and for DE analysis, are deposited to the NCBI Short Read Archive under accessions SRR15144741, SRR15144745, SRR15144747, and SRR15144751.

## Supporting Information

**Fig. S1** Leaf length and width phenotyping of the four selected accessions.

**Fig. S2** Differentially expressed transcripts and enriched GO terms in *O. glaberrima* and *O. australiensis*.

**Fig. S3** PCA-SOM clustering of gene expression across the selected rice accessions.

**Fig. S4** Quantification of mature fourth leaf length across the selected cultivated and wild rice accessions used for GA quantification.

**Fig. S5** Effects of GA and paclobutrazol treatments on leaf length and division zone size.

**Fig. S6** Cellular basis of leaf size differences across the selected rice accessions.

**Fig. S7** Expression analysis of *GRFs* and cell cycle-related genes in different growth zones of cultivated and wild rice accessions.

**Fig. S8** Expression analysis of GA- and cell cycle-related genes in response to GA and PAC treatment.

**Fig. S9** Silencing of positive control *OsChlH* and GA signaling repressor *OsSLR1*.

**Fig. S10** Expressions analysis of *GRFs* and *CYCLINs* in silencing lines of *OsGA20OX2 OsGRF7, OsGRF8*, and *OsSLR1*.

**Table S1** List of cultivated and wild rice accessions used in the present study

**Table S2** Quantification of maximum leaf width of fully-developed leaves of the seven cultivated and four wild rice accessions.

**Table S3** The total number of differentially expressed genes along with genes expressed at higher and lower levels for each pair-wise comparison.

**Table S4** Quantification of leaf kinematics parameters of the growing fourth leaf of *O. sativa* cv. Nipponbare under exogenous GA_3_ and paclobutrazol treatments.

**Table S5** Quantification of leaf kinematics parameters of the growing fourth leaf of *O. australiensis* under exogenous GA_3_ and paclobutrazol treatments.

**Table S6** List of primers used for gene expression analysis and transient silencing experiments.

**Dataset S1** List of statistically significant differentially expressed genes for each pair-wise comparison.

**Dataset S2** List of statistically significant GO-terms enriched for genes expressed at higher and lower levels in Fig. 2 and Supplemental Fig. S3.

**Dataset S3** List of genes present in each cluster derived from PCA-SOM analysis.

**Dataset S4** List of statistically significant GO-terms enriched for the genes present in the PCA-SOM clusters.

**Dataset S5** List of cell cycle-related genes, transcription factors and leaf developmental genes, and phytohormone-related genes present in clusters 1, 4, and 7 that were used for generating heatmaps in Fig. 2c-e and Fig. S3c.

**Dataset S6** List of genes present in the expression network shown in Fig. 2f with their network properties and module belongings.

**Dataset S7** Normalized read counts used for generating gene coexpression network.

