## Supplementary material for "Spatial control of cell division by GA-OsGRF7/8 module in a leaf explains the leaf length variation between cultivated and wild rice": Supplemntal Figures and Tables

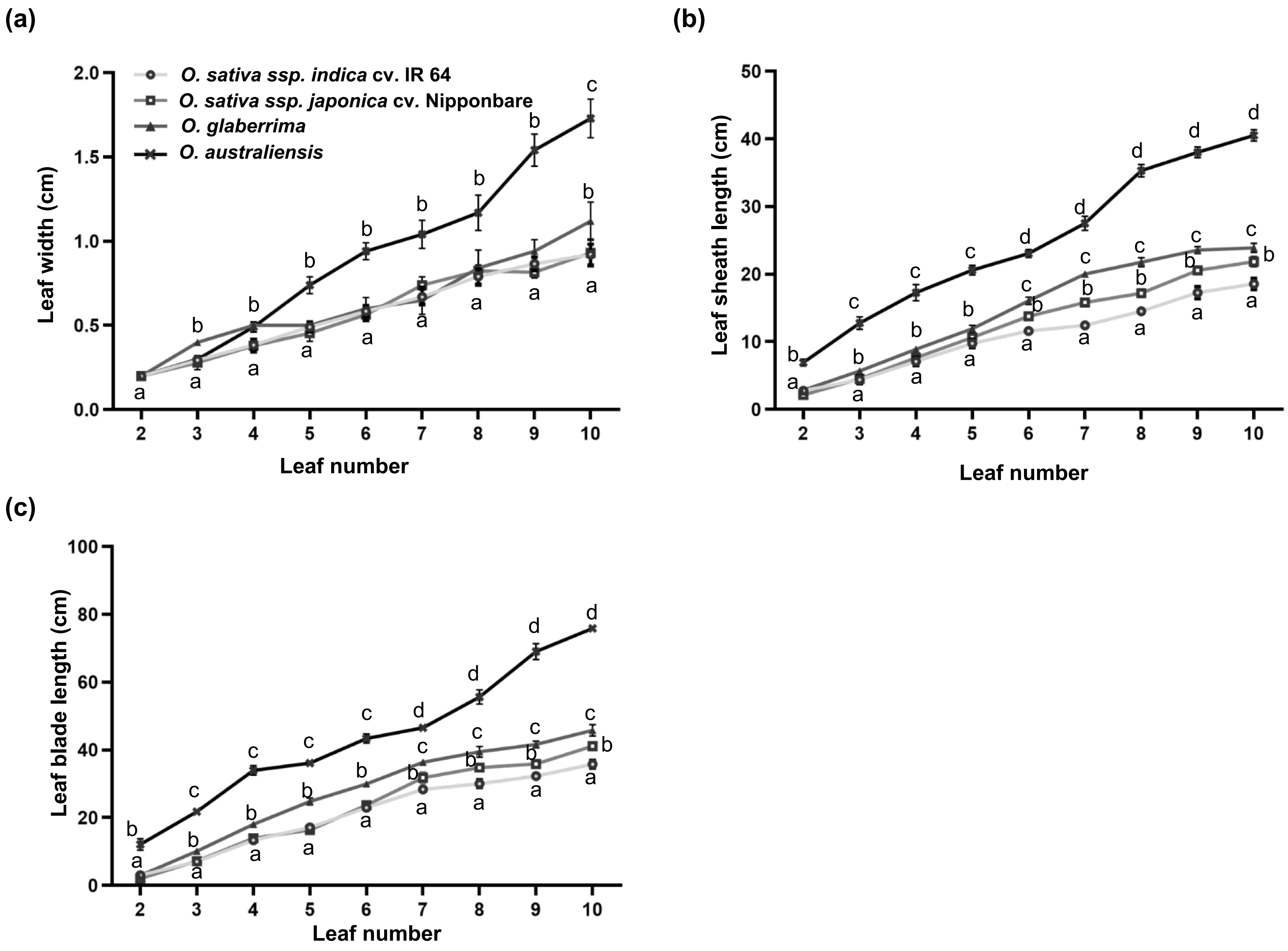


**Fig. S1** Leaf length and width phenotyping of the four selected accessions. Quantification of leaf blade width (a), leaf sheath length (b), and leaf blade length (c) of second leaf to tenth leaf of the selected rice accessions. Values in graphs represent mean±SD (n=15). Different letters indicate statistically significant differences according to one-way ANOVA followed by post-hoc Tukey HSD calculation at $P < 0.05$.

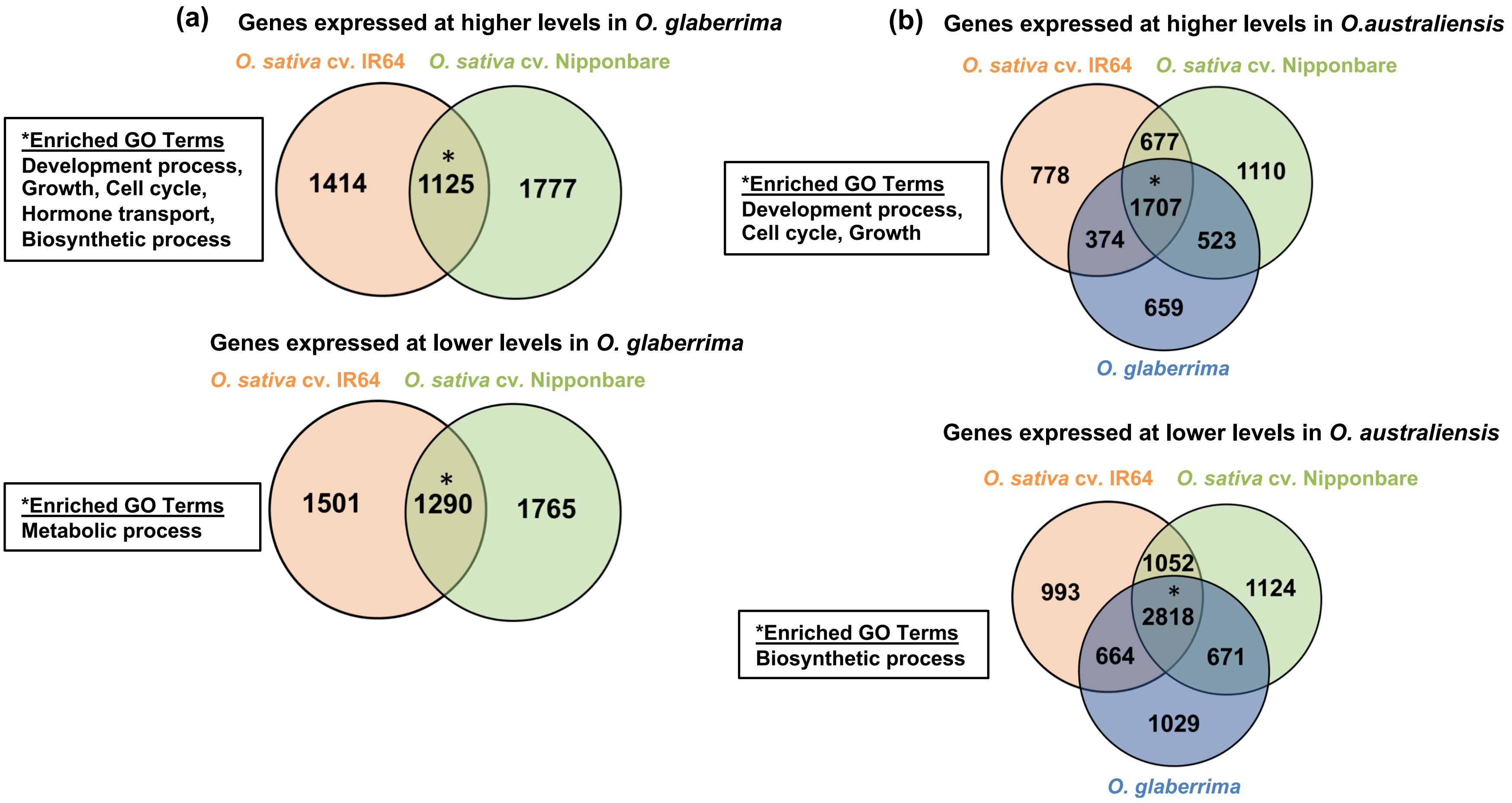


**Fig. S2 (**a) Differentially expressed transcripts and enriched GO terms in *O. glaberrima* compared to *O. sativa* cv. IR64 and *O. sativa* cv. Nipponbare. (b) Differentially expressed transcripts and enriched GO terms in *O. australiensis* compared to *O. sativa* cv. IR64, *O. sativa* cv. Nipponbare and *O. glaberrima*. * showed enriched GO terms.

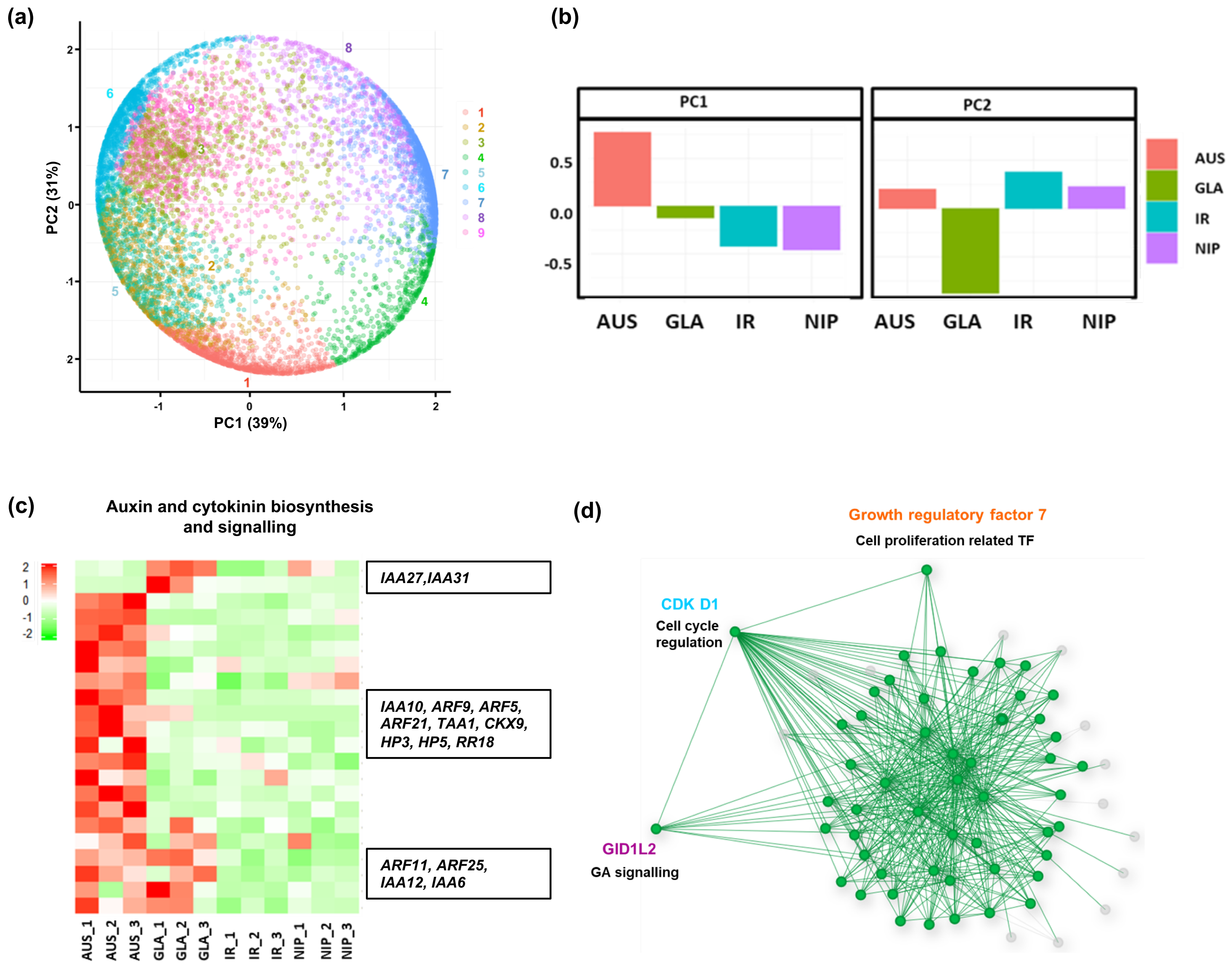


**Fig. S3** PCA-SOM clustering of gene expression across the selected rice accessions. (a) Expression profile of each transcript in PCA space with SOM node memberships represented by the different colours and numbers. (b) Loadings of principal components PC1 and PC2 with variance. (c) Expression patterns of auxin- and cytokinin-related genes in clusters 1, 4 and 7 across the four accessions. AUS= *O. australiensis*, GLA= *O. glaberrima*, IR= *O. sativa* cv. IR64, NIP= *O. sativa* cv. Nipponbare. (d) Connectivity in module 2 of gene regulatory network generated for the genes from the relevant PCA-SOM clusters.

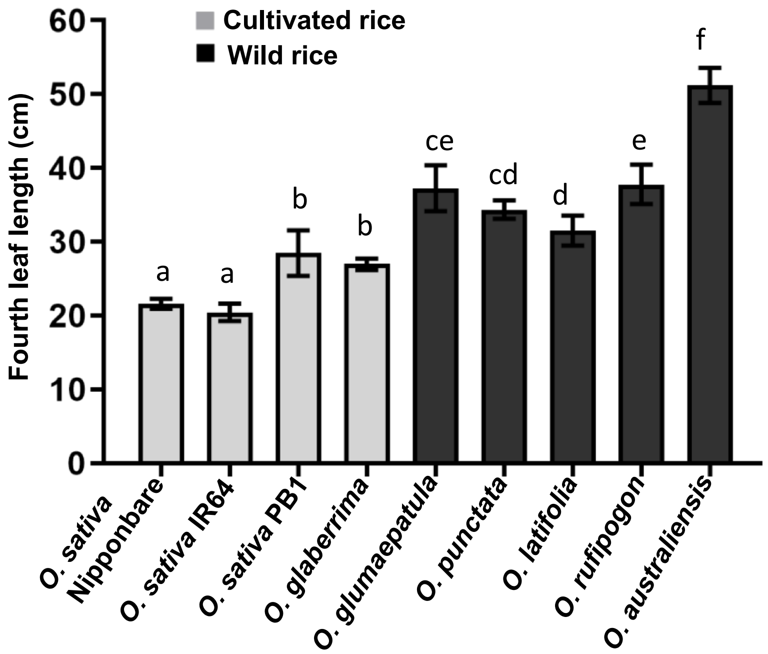


**Fig. S4** Quantification of mature fourth leaf length across the selected cultivated and wild rice accessions used for GA quantification. Values in graphs represent mean±SD (n=15). Different letters indicate statistically significant differences according to one-way ANOVA followed by post-hoc Tukey HSD calculation at $P < 0.05$.

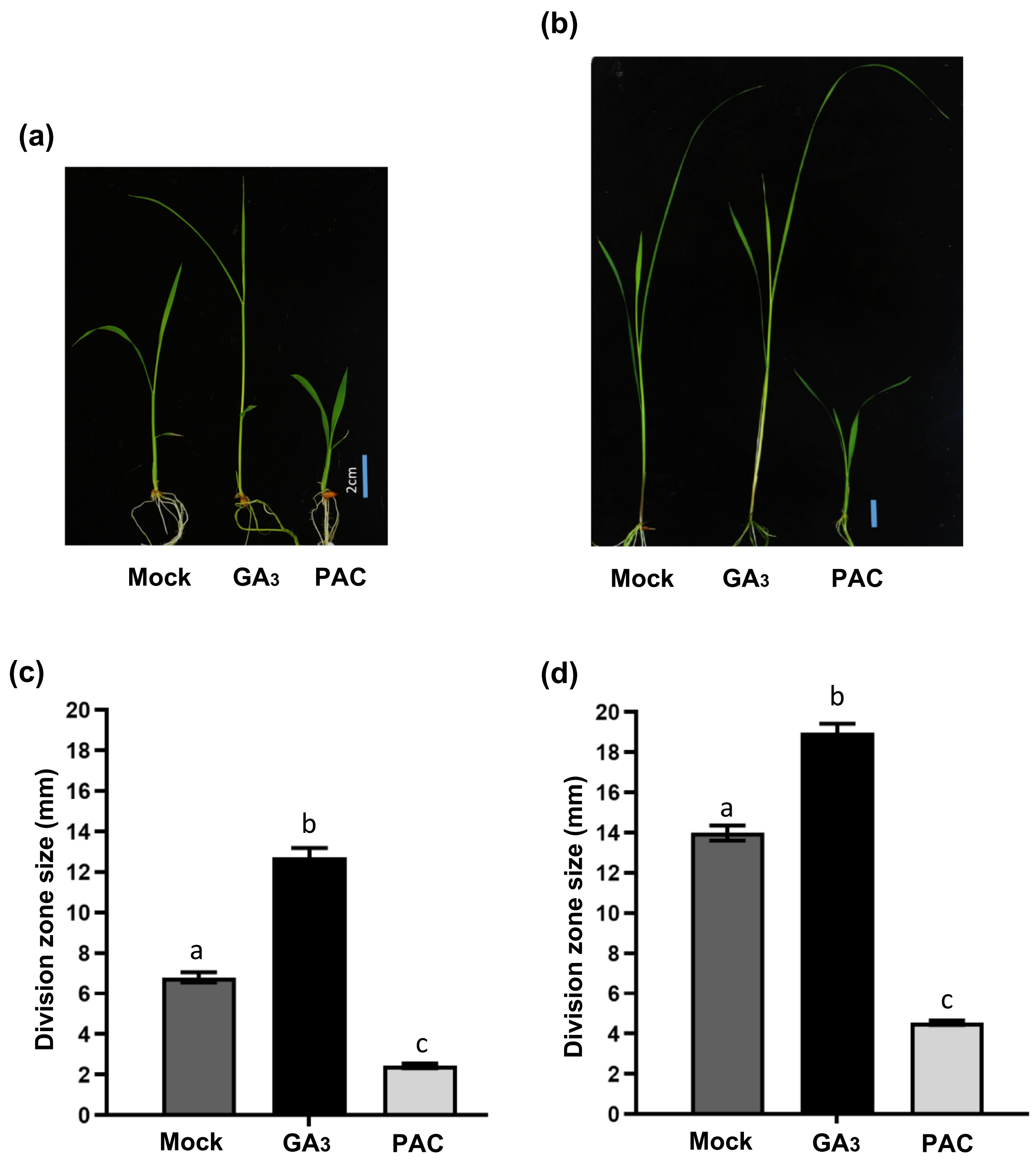


**Fig. S5** Effects of GA and paclobutrazol (PAC) treatments on leaf length and division zone size. (a, b) Effects of $GA_3$ (10µM) and PAC (1µM) treatments on leaf size and growth of *O. sativa* cv. Nipponbare (a) and *O. australiensis* (b). (c, d) $GA_3$ and PAC effects on division zone size of *O. sativa* cv. Nipponbare (c) and *O. australiensis* (d). Values represent mean ±SD (n=5). Different letters indicate statistically significant differences according to one-way ANOVA followed by post-hoc Tukey HSD calculation at $P < 0.05$.

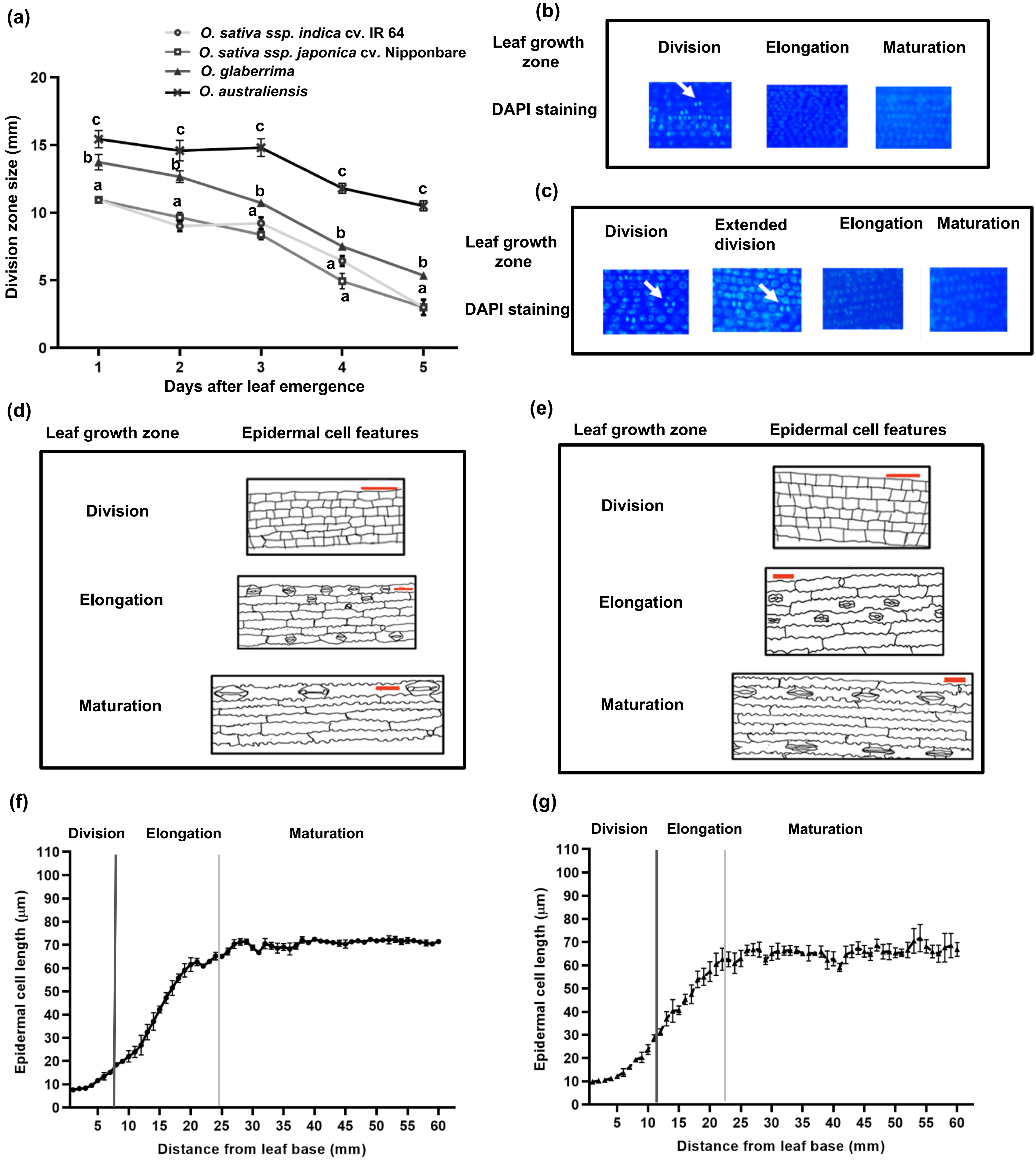


**Fig. S6** Cellular basis of leaf size differences across the selected rice accessions. (a) Quantification of division zone size of leaf four at different days of emergence for the selected rice accessions. Values show mean±SD (n=5). Different letters indicate statistically significant differences according to one-way ANOVA followed by post-hoc Tukey HSD calculation at P < 0.05. (b, c) Representative DAPI staining images for the different leaf-growth-zones of *O. sativa* cv. Nipponbare (b) and *O. australiensis* (c). White arrows show DAPI-stained nuclei indicating dividing cells. (d, e) Representative hand-drawn microscopic images of epidermal cells for the different leaf-growth-zones of *O. sativa* cv. Nipponbare (d) and *O. australiensis* (e). Scale bar = 20µm. (f, g), Different growth zones on fourth leaf of *O. sativa* cv. IR64 (f) and *O. glaberrima* (g) at second day of emergence as quantified by epidermal cell length profiling and DAPI staining. Values represent mean ±SD (n=5). Vertical lines on graph shows different growth zones.

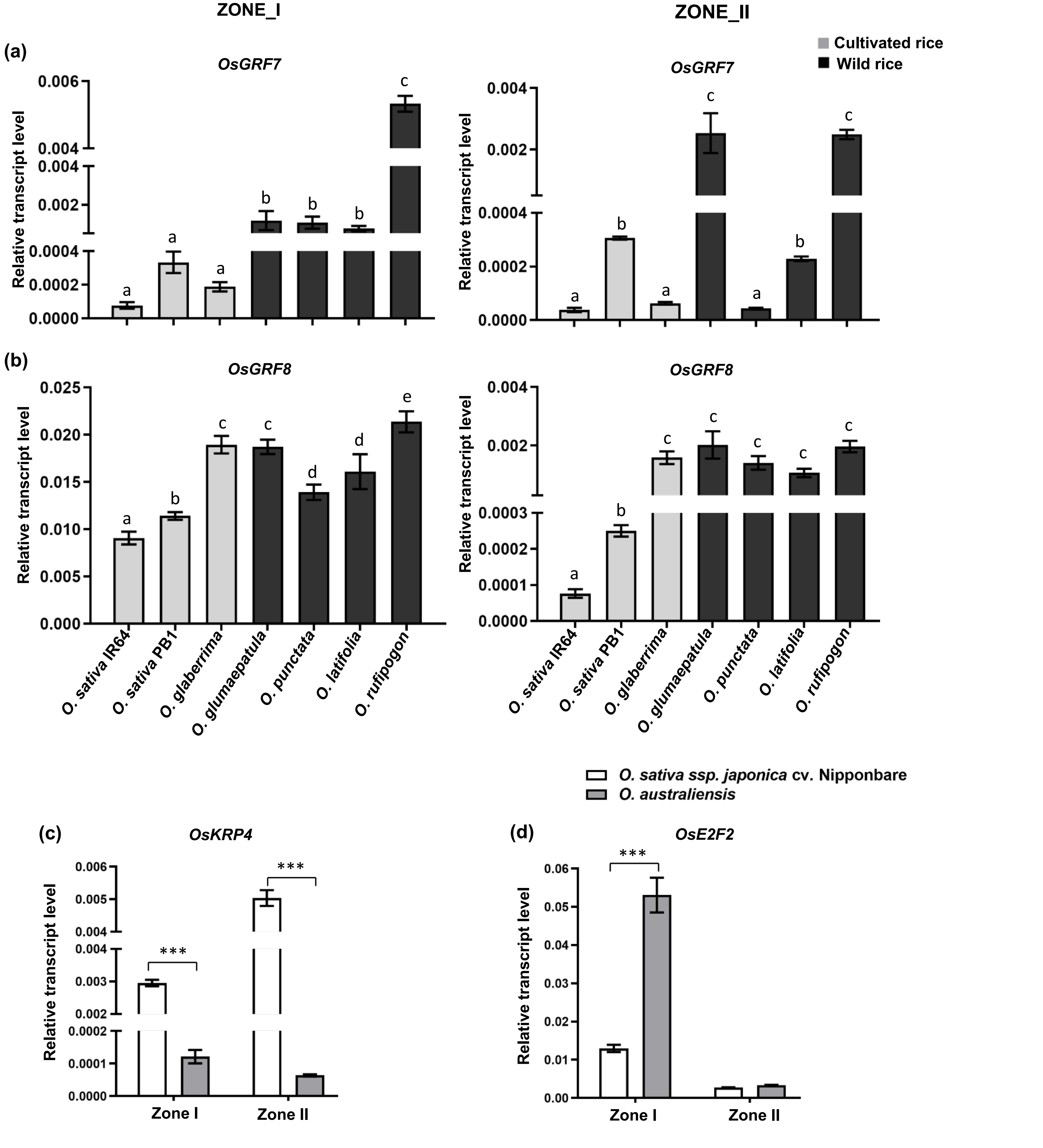


**Fig. S7** Expression analysis of *GRFs* and cell cycle-related genes in different growth zones of cultivated and wild rice accessions. (a, b) Expression analysis of *OsGRF7* (a) and *OsGRF8* (b) across the selected cultivated and wild rice accessions. (c, d) Expression analysis of *OsKRP4* (c) and *OsE2F2* (d) in the two growth zones of *O. sativa* cv. Nipponbare and *O. australiensis.* Zone I represents region from leaf base to 0.8cm and zone II represents 0.8cm to 1.6cm from leaf base. Data shown are mean ±SD of three biological replicates, *** $P < 0.001$ by Student's t-test.

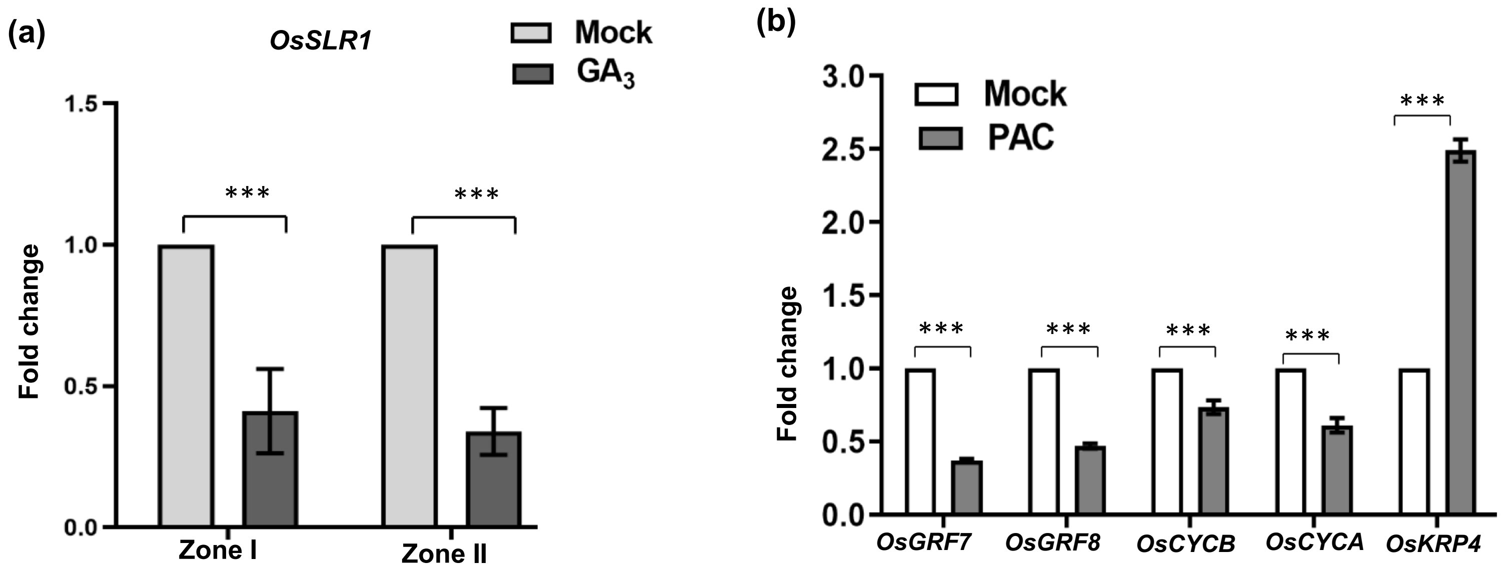


**Fig. S8** Expression analysis of GA- and cell cycle- related genes in response to GA and PAC treatment. (a) Expression analysis of *OsSLR1* in response to $GA_3$ treatment in the two growth zones of *O. sativa* cv. Nipponbare. Zone I represents region from leaf base to 0.8cm and zone II represents 0.8cm to 1.6cm from leaf base. (b) Effect of PAC treatment on expression of *GRF*s and cell-cycle-related genes in the division zone of *O. sativa* cv. Nipponbare. Data shown are mean ±SD of three biological replicates, *** $P < 0.001$ by Student's t-test.

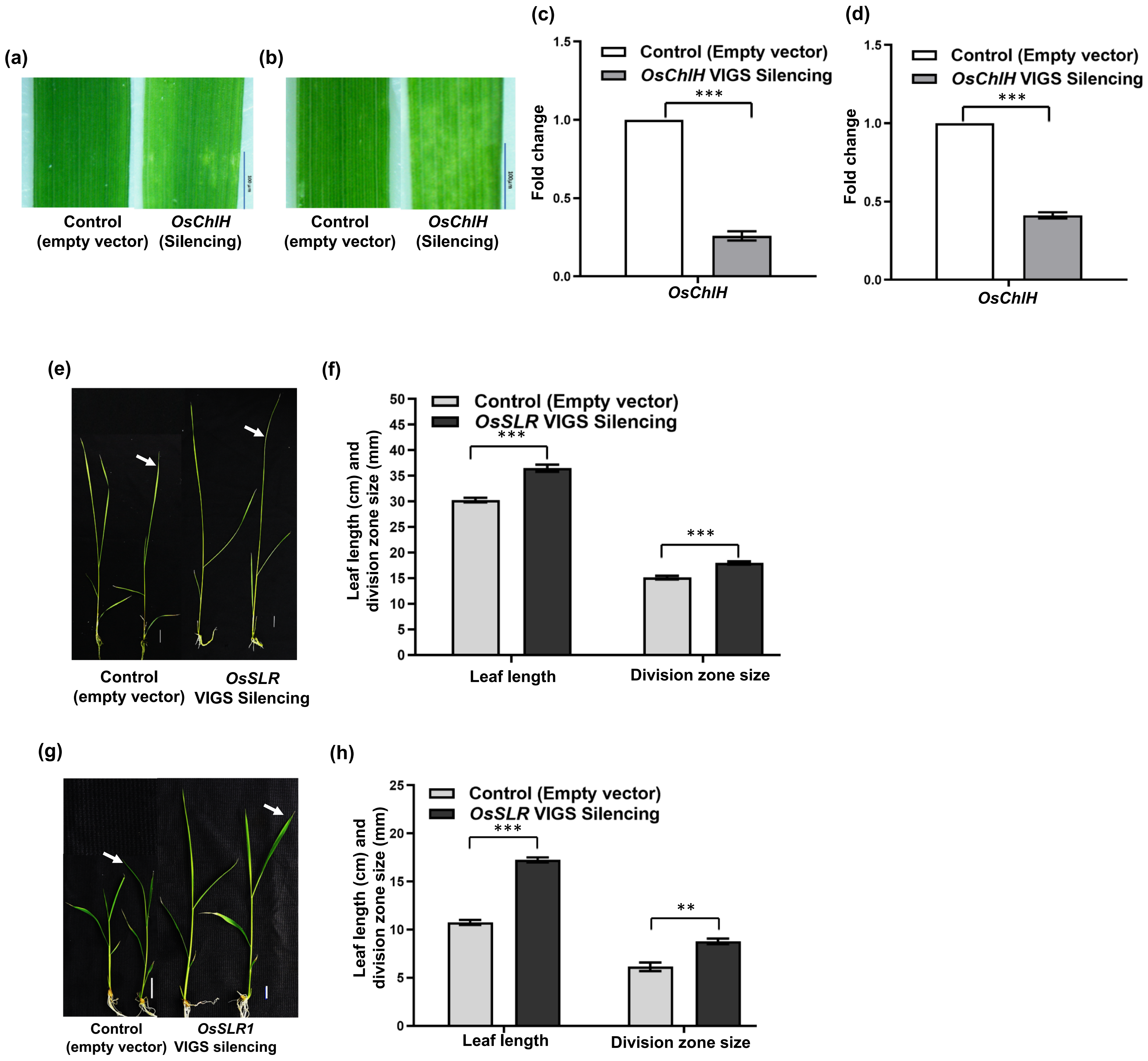


**Fig. S9** Silencing of positive control *OsChlH* and GA signalling repressor *OsSLR1*. (a, b) Phenotype of *OsChlH* silencing line of *O. sativa* cv. Nipponbare along with control (a) and *O. australiensis* along with control (b). (c, d) Expression levels of *ChlH* gene in the *OsChlH* silencing line and control plants of *O. sativa* cv. Nipponbare (c) and *O. australiensis* (d). (e) Shown are the leaf phenotypes of *OsSLR1* silencing in *O. australiensis* along with the control plants. (f) Effect of *OsSLR1* silencing on leaf length and size of division zone in *O. australiensis*. (g) Shown are the leaf phenotypes of *OsSLR1* silencing in *O. sativa* cv. Nipponbare along with the control plants. (h) Effect of *OsSLR1* silencing on leaf length and size of division zone in *O. sativa* cv. Nipponbare. Data represent mean ±SD of at least three biological replicates, *** $P < 0.001$ and ** $P < 0.01$ by Student's t-test. White arrow indicates fourth leaf and scale bar = 2cm in (e) and (g).

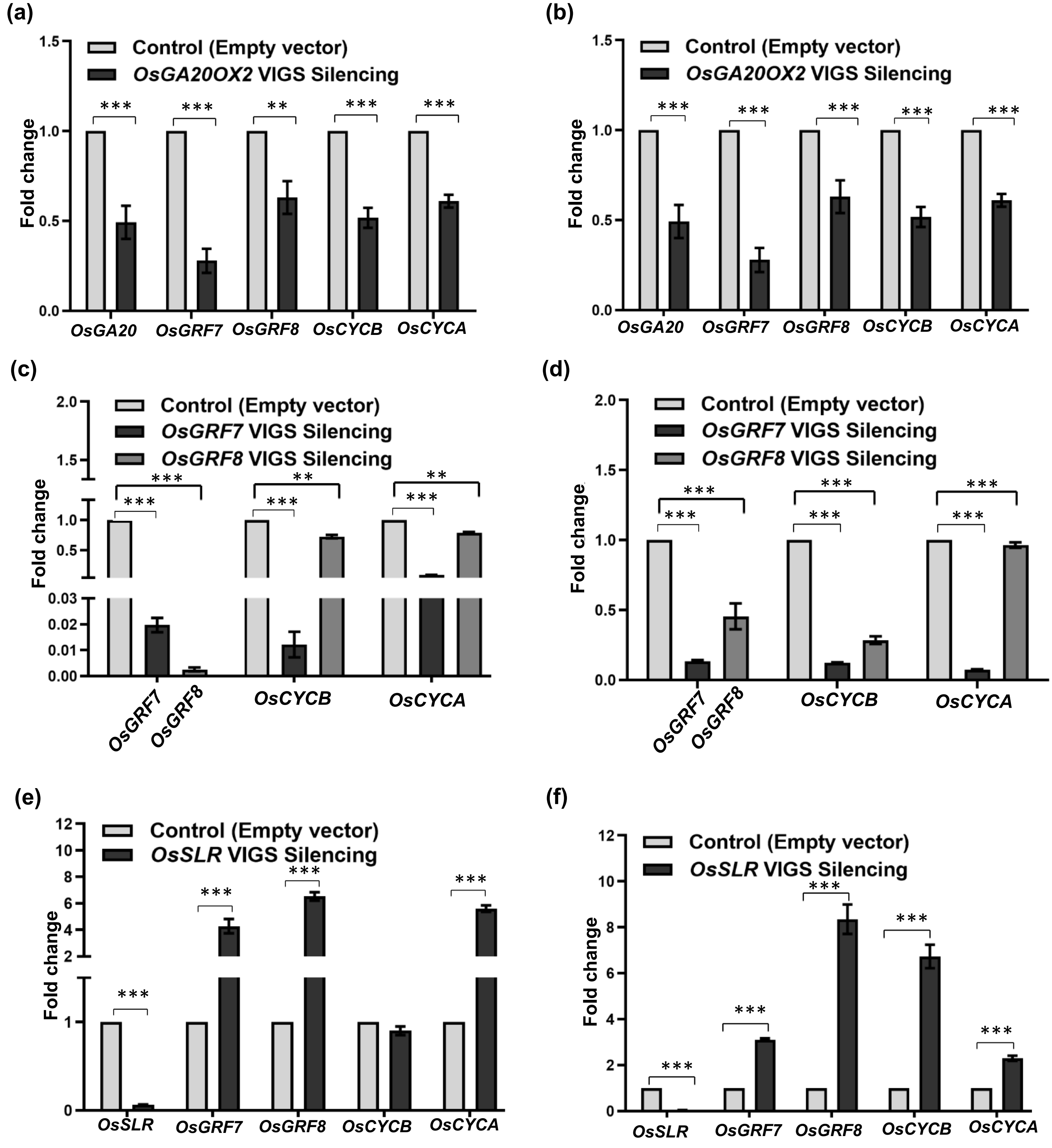


**Fig. S10** Expressions analysis of *GRFs* and *CYCLINs* in silencing lines of *OsGA20OX2 OsGRF7, OsGRF8,* and *OsSLR1* in *O. australiensis* and *O. sativa* cv. Nipponbare. (a, b) Expression levels of target GA biosynthetic gene, *GRFs*, and *CYCLINs* in *OsGA20OX2* silencing lines of *O. australiensis* (a) and *O. sativa* cv. Nipponbare (b). (c, d) Expression analysis of target genes and *CYCLINs* in *OsGRF7* and *OsGRF8* silencing lines of *O. australiensis* (c) and *O. sativa* cv. Nipponbare (d). (e-f) Expression analysis of target *OsSLR1*, *GRFs,* and *CYCLINs* in *OsSLR1* silencing lines of *O. australiensis* (e) and *O. sativa* cv. Nipponbare (f). Data represent mean ±SD of at least three biological replicates, *** $P < 0.001$ by Student's t-test.

**Table S1** List of cultivated and wild rice accessions used in the present study.

| Oryza species | Genome | Cultivated/ Wild | Cultivar/ accession number |
|---|---|---|---|
| *Oryza sativa ssp. japonica* | AA | Cultivated | Nipponbare |
| *Oryza sativa ssp. indica* | AA | Cultivated | IR 64 |
| *Oryza sativa ssp. indica* | AA | Cultivated | Vandana |
| *Oryza sativa ssp. indica* | AA | Cultivated | Swarna |
| *Oryza sativa ssp. indica* | AA | Cultivated | Pusa Basmati1 |
| *Oryza sativa ssp. indica* | AA | Cultivated | Nagina 22 |
| *Oryza glaberrima* | AA | Cultivated | IR 102925 |
| *Oryza glumaepatula* | AA | Wild | IR104151 |
| *Oryza punctata* | BB | Wild | IRGC105137 |
| *Oryza latifolia* | CCDD | Wild | IRGC 99596 |
| *Oryza rufipogon* | AA | Wild | IRGC 99562 |
| *Oryza australiensis* | EE | Wild | IRGC 105272 |

**Table S2** Quantification of maximum leaf width (cm) of fully-developed second leaf to the eighth leaf of the seven cultivated and five wild rice accessions. Each value represents mean ± SD, where n = 15 data points from different plants. Different letters indicate statistically significant differences according to one-way ANOVA followed by post-hoc Tukey HSD calculation at P < 0.05.

| **Oryza species** | **Variety/ accession number** | **Leaf number** | | | | | | |
|---|---|---|---|---|---|---|---|---|
| | | **Leaf 2** | **Leaf 3** | **Leaf 4** | **Leaf 5** | **Leaf 6** | **Leaf 7** | **Leaf 8** |
| *Oryza sativa ssp. japonica* | Nipponbare | 0.20 ± 0.01 (a) | 0.28 ± 0.05 (ac) | 0.38 ± 0.04 (a) | 0.46 ± 0.05 (ad) | 0.57 ± 0.05 (ac) | 0.74 ± 0.05 (acd) | 0.83 ± 0.07 (acf) |
| *Oryza sativa ssp. indica* | IR 64 | 0.20 ± 0.001 (a) | 0.30 ± 0.02 (c) | 0.39 ± 0.03 (a) | 0.49 ± 0.03 (ae) | 0.59 ± 0.03(ac) | 0.67 ± 0.05 (abcd) | 0.79 ± 0.05 (abe) |
| *Oryza sativa ssp. indica* | Vandana | 0.20 ± 0.001 (a) | 0.23 ± 0.05 (ad) | 0.29 ± 0.03 (b) | 0.37 ± 0.05 (b) | 0.46 ± 0.05(b) | 0.58 ± 0.07 (b) | 0.69 ± 0.06 (b) |
| *Oryza sativa ssp. indica* | Swarna | 0.21 ± 0.03 (a) | 0.32 ± 0.04 (c) | 0.40 ± 0.001 (a) | 0.43 ± 0.05 (be) | 0.59 ± 0.06 (ac) | 0.72 ± 0.09 (ce) | 0.85 ± 0.07 (ce) |
| *Oryza sativa ssp. indica* | Pusa Basmati1 | 0.21 ± 0.03 (a) | 0.31 ± 0.03 (c) | 0.35 ± 0.05 (ab) | 0.48 ± 0.04 (a) | 0.64 ± 0.05 (cd) | 0.77 ± 0.04 (c) | 0.89 ± 0.06 (c) |
| *Oryza sativa ssp. indica* | Nagina 22 | 0.20 ± 0.001 (a) | 0.31 ± 0.03 (c) | 0.33 ± 0.04 (ab) | 0.42 ± 0.03 (bcd) | 0.55 ± 0.05 (abc) | 0.77 ± 0.05 (c) | 1.03 ± 0.05 (d) |
| *Oryza glaberrima* | IR 102925 | 0.20 ± 0.001 (a) | 0.40 ± 0.001 (b) | 0.50 ± 0.001 (c) | 0.50 ± 0.001 (a) | 0.60 ± 0.07 (ad) | 0.65 ± 0.08 (bde) | 0.84 ± 0.11 (ce) |
| *Oryza glumaepatula* | IR104151 | 0.20 ± 0.001 (a) | 0.30 ± 0.05 (c) | 0.34 ± 0.05 (ab) | 0.46 ± 0.05 (a) | 0.62 ± 0.03 (ad) | 0.68 ± 0.04 (bde) | 0.81 ± 0.03 (ce) |
| *Oryza punctata* | IRGC105137 | 0.20 ± 0.001 (a) | 0.31 ± 0.03 (c) | 0.31 ± 0.03 (b) | 0.40 ± 0.01 (bd) | 0.54 ± 0.05 (ab) | 0.64 ± 0.05 (bde) | 0.79 ± 0.03 (fe) |
| *Oryza latifolia* | IRGC 99596 | 0.31 ± 0.03 (b) | 0.41 ± 0.03 (b) | 0.53 ± 0.05 (c) | 0.83 ± 0.07 (f) | 1.30 ± 0.16 (e) | 1.80 ± 0.22 (f) | 1.94 ± 0.12 (g) |
| *Oryza rufipogon* | IRGC 99562 | 0.31 ± 0.02(b) | 0.39 ± 0.03(b) | 0.60 ± 0.001(d) | 0.67 ± 0.02(g) | 0.88 ± 0.02(f) | 1.08 ± 0.01(g) | 1.35 ± 0.1 (h) |
| *Oryza australiensis* | IRGC 105272 | 0.20 ± 0.001 (a) | 0.30 ± 0.001 (c) | 0.49 ± 0.03 (c) | 0.74 ± 0.05 () | 0.94 ± 0.05 (f) | 1.04 ± 0.08 (g) | 1.06 ± 0.12 (d) |

**Table S3** The total number of differentially expressed genes along with genes expressed at higher and lower levels for each pair-wise comparison

| Species Comparisons | Differentially expressed genes | Genes expressed at higher levels | Genes expressed at lower levels |
|---|---|---|---|
| *O. sativa* cv. IR64 Vs *O. sativa* cv. Nipponbare | 5414 | 2717 | 2697 |
| *O. glaberrima* Vs *O. sativa* cv. IR64 | 5330 | 2539 | 2791 |
| *O. australiensis* Vs *O. sativa* cv. IR64 | 9063 | 3536 | 5527 |
| *O. glaberrima* Vs *O. sativa* cv. Nipponbare | 5957 | 2902 | 3055 |
| *O. australiensis* Vs *O. sativa* cv. Nipponbare | 9782 | 4017 | 5765 |
| *O. australiensis* Vs *O. glaberrima* | 8445 | 3263 | 5182 |

**Table S4** Quantification of leaf kinematics parameters of the growing fourth leaf of *O. sativa* cv. Nipponbare under exogenous $GA_3$ (10µM) and paclobutrazol (1 µM) treatments along with the control (Mock) plants. Each value represents mean ± SD (n = 15 for mature leaf length and leaf elongation rate, n = 5 for all other parameters). Different letters indicate statistically significant differences according to one-way ANOVA followed by post-hoc Tukey HSD calculation at $P < 0.05$.

| **Growth parameters** | **Mock** | **$GA_3$** | **PAC** |
|---|---|---|---|
| Mature leaf length (cm) | 22.02 ± 1.42(a) | 35.82 ± 1.06(b) | 11.07 ± 0.46(c) |
| Leaf Elongation Rate (mm/h) | 1.41 ± 0.01(a) | 2.78 ± 0.03(b) | 0.89 ± 0.01(c) |
| Meristem length (mm) | 6.45 ± 0.36(a) | 12.72 ± 0.49(b) | 2.60 ± 0.18(c) |
| Length of the growth zone (mm) | 33.00 ± 1.00(a) | 34.33 ± 1.00(a) | 24.67 ± 0.58(b) |
| Mature cell length (µm) | 64.71 ± 0.58(a) | 62.37 ± 0.15(ab) | 58.55 ± 0.97(b) |
| Cell production rate (cells/h) | 21.81 ± 0.24(a) | 44.72 ± 0.09(b) | 15.22 ± 0.24(c) |
| Number of cells in the meristem | 616.80 ± 23.90(a) | 1071.10 ± 72.10(b) | 306.20 ± 36.80(c) |
| Number of cells in the growth zone | 1337.60 ± 41.40(a) | 1788.10 ± 69.70(b) | 1285.30 ± 51.40(a) |
| Number of cells in the elongation zone | 720.80 ± 36.50(a) | 716.90 ± 29.07(a) | 979.08 ± 14.60(b) |
| Average cell division rate (cell. $cell^{-1}.h^{-1}$) | 0.04 ± 0.001(a) | 0.04 ± 0.001(a) | 0.05 ± 0.001(b) |
| Cell cycle duration (h) | 19.61 ± 0.97(a) | 16.62 ± 1.13(b) | 13.94 ± 1.45(c) |
| Time in elongation zone (h) | 33.05 ± 1.70(a) | 16.14 ± 0.63(b) | 64.34 ± 0.08(c) |
| Time in division zone (h) | 181.70 ± 10.08(a) | 157.40 ± 12.30(b) | 125.90 ± 12.50(c) |
| Length of cells leaving meristem (µm) | 14.90 ± 0.16(a) | 21.29 ± 3.36(b) | 7.61 ± 1.68(c) |
| Average cell expansion rates (µm. $µm^{-1}.h^{-1}$) | 0.04 ± 0.002(a) | 0.07 ± 0.001(b) | 0.03 ± 0.003(c) |

**Table S5** Quantification of leaf kinematics parameters of the growing fourth leaf of *O. australiensis* under exogenous $GA_3$ (10µM) and paclobutrazol (1 µM) treatments along with the control (Mock) plants. Each value represents mean ± SD (n = 15 for mature leaf length and leaf elongation rate, n = 5 for all other parameters). Different letters indicate statistically significant differences according to one-way ANOVA followed by post-hoc Tukey HSD calculation at $P < 0.05$.

| **Growth parameters** | **Mock** | **$GA_3$** | **PAC** |
|---|---|---|---|
| Mature leaf length (cm) | 43.80± 3.15(a) | 60.75 ± 4.66(b) | 17.12 ± 2.29(c) |
| Leaf Elongation Rate (mm/h) | 2.31 ± 0.01(a) | 4.11 ± 0.08(b) | 1.70 ± 0.01(c) |
| Meristem length (mm) | 14.15 ± 0.43(a) | 19.13 ± 0.18(b) | 5.00 ± 1.00(c) |
| Length of the growth zone (mm) | 34.67 ± 0.58(a) | 33.33 ± 1.15(a) | 25.60 ± 1.15(b) |
| Mature cell length (µm) | 80.36 ± 0.73(a) | 78.28 ± 0.49(b) | 83.09 ± 0.49(c) |
| Cell production rate (cells/h) | 28.73 ± 0.32(a) | 52.49 ± 1.28(b) | 20.51 ± 0.17(c) |
| Number of cells in the meristem | 943.00 ± 20.07(a) | 1223.07 ± 51.10(b) | 493.80 ± 65.20(c) |
| Number of cells in the growth zone | 1361.50 ± 60.20(a) | 1532.10 ± 43.50(a) | 998.80 ± 41.60(b) |
| Number of cells in the elongation zone | 418.50 ± 45.10(ab) | 309.06 ± 20.50(b) | 505.00 ± 105.01(ac) |
| Average cell division rate (cell. $cell^{-1}.h^{-1}$) | 0.03 ± 0.001(a) | 0.04 ± 0.001(b) | 0.04 ± 0.001(b) |
| Cell cycle duration (h) | 22.70 ± 0.40(a) | 16.10 ± 0.30(b) | 16.60 ± 2.08(b) |
| Time in elongation zone (h) | 14.50 ± 1.40(a) | 5.80 ± 0.50(b) | 24.60 ± 5.30(c) |
| Time in division zone (h) | 224.80 ± 4.80(a) | 154.10 ± 16.70(b) | 152.70 ± 16.10(b) |
| Length of cells leaving meristem (µm) | 31.50 ± 2.50(a) | 38.20 ± 1.20(b) | 14.10 ± 2.20(c) |
| Average cell expansion rates (µm. $µm^{-1}.h^{-1}$) | 0.06 ± 0.01(a) | 0.12 ± 0.02(b) | 0.07 ± 0.01(a) |

**Table S6** List of primers used for gene expression analysis and transient silencing experiments.

| Acronym | Forward (5´-3´) | Reverse (5´-3´) |
|---|---|---|
| **qRT-PCR analysis** | | |
| *GA20OX2* | GCGAGGAGATGAAGGAGCTG | ATGGCGGGTAGTAGTTGCAC |
| *GA3OX2* | ACTCGGGCTTCTTCACCTTC | CCGTTGGTGAGGATGTGGAA |
| *GA2OX4* | GATCACATCCCGCTGCTGAG | TGACGACCTTGAAGAACCCG |
| *SLR1* | CGCTGCACTACTACTCCACC | GGTACACCTCGGACATGACC |
| *GRF7* | GAAACAACACGCGAGCAAGT | AGAGGTTCAAAGATGGCGGG |
| *GRF8* | TTCTGTCCCTTCCAGCTTGC | GGCATCTTCTGGGTTCTACA |
| *CYCB1;4* | GCTCGAGGAAGAAGGTCATC | CGAGCTTGTCAATGTCCTCA |
| *CYCA3;2* | TACGGCTGTGTTCGGCTC | CCTTCAACTCAGATGCCCTG |
| *KRP4* | GCAGCCACTCGAGTTCTCAT | GATGAAAGCCTGTCGTTGCC |
| *E2F2* | AGTGGCATGGAGACTCCTC | TTGGCTGTGACTCTGCAGC |
| *ChlH* | GCTATGTTGAACCTGGCCCT | CAAGGCAGCTGTAGTTGGGA |
| *ACTIN* | GAAGTGCGACGTGGATATTAG | CAGACACTGTACTTCCTTTCAG |
| **Gene-silencing** | | |
| *GA20OX2* | TTAATTAAACGGGTTCTTCCAGGTGTC | ACGCGTCGCGAAGAACTCCCTGTAGT |
| *SLR1* | TTAATTAATCGTCACCGTGGTAGAGCAG | ACGCGTCTCGCCTGTTTGTAGGCATT |
| *GRF7* | TTAATTAATCCTCCTCAAGCCCAGGATT | ACGCGTTGGTGTGAATGGCCCCCTAA |
| *GRF8* | TTAATTAAATCATGAGCTCAATGCCACA | ACGCGT CCCATCTTGCTCTTGCTCTT |
| *ChlH* | TTAATTAAGCCACCAAGTGCCATCGA | ACGCGTCTCTCTGTCTGCTCTGA |
